# Nucleosome remodeling by a CHD enzyme promotes H3K9 methylation establishment and spreading via remodeler-writer feedback

**DOI:** 10.64898/2026.05.03.722496

**Authors:** Melissa Seman, Alexandria Latuda, Agnisrota Mazumder, Luca Marco Wolfstädter, Fengting Huang, Amith Zafal Abdulla, Sigurd Braun, Kaushik Ragunathan

## Abstract

In *Schizosaccharomyces pombe*, the conserved CHD remodeler Mit1 function within the SHREC remodeler-deacetylase complex (a homolog of the metazoan Mi-2/NuRD complex), which is essential for H3K9 methylation-dependent heterochromatin establishment. However, the mechanism by which remodeler activity promotes silencing is unknown. Current models posit a hierarchical relationship between histone modifications and remodeler activity, with Mit1 acting exclusively downstream of H3K9 methylation. Here, we challenge this model by showing that tethering Mit1 at an ectopic site within euchromatin is sufficient to initiate heterochromatin assembly and generate extended domains of *de novo* H3K9 methylation. This process requires the Mit1 catalytic activity but does not involve direct physical interaction with Clr4, suggesting Mit1-mediated nucleosome remodeling creates a chromatin context that enhances Clr4 function. Using a genome-wide deletion screen, we determined that Mit1-initiated silencing requires all core heterochromatin factors and is critically dependent on Clr4 dosage. Furthermore, Mit1 activity facilitates heterochromatin spreading at subtelomeric regions and promotes H3K9 methylation at novel genomic sites implicated in cellular adaptation. Together, our findings support a model in which remodeler-writer pairs, analogous to reader-writer pairs, constitute conserved regulatory modules through which nucleosome organization directs the establishment of heritable epigenetic states.

## INTRODUCTION

Genome organization is dynamic and subject to constant change thus enabling eukaryotic cells to balance the inherent tradeoffs between DNA accessibility and the need for compaction (Wu et al., 2009). These opposing conformational states represent two ends of a regulatory spectrum that governs gene expression. Actively transcribed regions of the genome adopt an accessible conformation that facilitates RNA polymerase II activity, whereas silent regions of the genome adopt conformational states that suppress transcription (Jiang and Pugh, 2009; Li et al., 2007; Swygert and Peterson, 2014). Nucleosome occupancy is central to this regulatory logic (Radman-Livaja and Rando, 2010; Segal and Widom, 2009). Nucleosomes positioned across regulatory sequences can restrict transcription factor binding, while their removal exposes the underlying DNA sequence to the transcriptional machinery. How nucleosomes are organized directly impacts whether a locus is transcriptionally active or silent.

The structural plasticity of chromatin depends on the enzymatic activity of ATP-dependent chromatin remodelers (Tsukiyama, 2002). These enzymes use ATP hydrolysis to slide, evict, or exchange nucleosomes, thereby altering chromatin organization genome-wide (Clapier et al., 2017; Zhou et al., 2016). The contribution of chromatin remodelers to transcription is well established. SWI/SNF-family remodelers disrupt or evict nucleosomes at promoters and enhancers to generate accessible chromatin for transcription factor binding (Ahmad et al., 2024; Côté et al., 1994; Kwon et al., 1994; Tsukiyama et al., 1994). CHD- and ISWI-family remodelers alter chromatin organization through nucleosome sliding at promoters and within transcribed genes (Smolle et al., 2012; Whitehouse et al., 2007; Zentner et al., 2013). The overarching principle that emerges from these findings is that remodelers alter DNA exposure, thereby regulating access to the transcriptional machinery. This suggests a relatively straightforward logic for how remodelers activate gene expression. In contrast, how remodelers contribute to transcriptional silencing especially within regions such as constitutive heterochromatin remains far less understood.

Chromatin remodelers are an integral part of silencing complexes and are broadly required for heterochromatin establishment across eukaryotes. The **<u>Nu</u>**cleosome **<u>R</u>**emodeling and **<u>D</u>**eacetylase complex (Mi-2/NuRD), which couples the nucleosome sliding activity of CHD3/CHD4 remodelers with the deacetylase activity of HDAC1/2, is important for silencing developmental genes in metazoans (Bornelöv et al., 2018; Low et al., 2020; Tong et al., 1998; Xue et al., 1998; Zhang et al., 1999). CHD-family remodelers are often part of silencing complexes that consist of histone-modifying enzymes or chromatin readers. In embryonic stem cells, the ChAHP complex pairs CHD4 with HP1 proteins and the DNA-binding factor ADNP to transcriptionally silence SINE elements (Ostapcuk et al., 2018). Yet, the functional significance of remodelers and their role in silencing gene expression remains unclear.

One possible explanation is that these remodelers enforce silencing by simply increasing nucleosome density and eliminating nucleosome-free regions within heterochromatin (Bornelöv et al., 2018; Whitehouse et al., 2007). However, nucleosome occupancy within heterochromatin is comparable to the rest of the genome and is instead characterized by irregularly spaced nucleosomes as opposed to being compact(Abdulhay et al., 2020; Garcia et al., 2010; Kerr et al., 2023; Marr et al., 2021; Prajapati et al., 2025). Hence, the contribution of remodelers to silencing must extend beyond simply restricting DNA accessibility, pointing to other mechanisms that likely connect remodeler activity to the establishment of silent epigenetic states.

A system in which a single remodeler is genetically essential for heterochromatin establishment provides a unique opportunity to address this question. In the model organism *Schizosaccharomyces pombe*, the CHD-family chromatin remodeler, Mit1, is essential for heterochromatin establishment and is part of a remodeler-deacetylase complex called SHREC which has striking parallels to the NuRD complex in metazoans (Creamer et al., 2014; Job et al., 2016; Motamedi et al., 2008; Sugiyama et al., 2007). Deleting Mit1 disrupts transcriptional silencing across all constitutive heterochromatin loci (pericentromeres, telomeres, and the mating type locus) clearly underscoring the essential role of chromatin remodeling in heterochromatin assembly (Creamer et al., 2014). The primary function of Mit1 is thought to involve changes in the distribution of nucleosome free regions given its ability *in vitro* to slide centered nucleosomes to the ends of a DNA fragment (Creamer et al., 2014; Garcia et al., 2010). Since Mit1 is recruited to heterochromatin regions by interacting with the HP1 protein Chp2, our current understanding is that Mit1 functions solely downstream of H3K9me with no role during the initiation process (Leopold et al., 2019a; Maksimov et al., 2018). To understand how Mit1 associated remodeling activity affects transcriptional silencing, we used a synthetic approach to tether Mit1 at an ectopic site (Audergon et al., 2015; Donovan et al., 2019; Ni and Muegge, 2021; Ragunathan et al., 2015; Scacchetti et al., 2018). Using this approach, our goal was to isolate how Mit1 remodeling activity influences transcriptional silencing.

Remarkably, we found that tethering Mit1 with intact ATPase activity can establish transcriptional silencing and nucleate an epigenetic state characterized by H3K9me. This observation suggests that Mit1 remodeling activity can promote H3K9me establishment, challenging the canonical view of remodelers as solely acting downstream of histone modifications. Instead, our results imply that Mit1 can remodel nucleosomes to adopt configurations that can be exploited by enzymes such as Clr4 to establish transcriptional silencing. More broadly, our findings reveal a previously unrecognized role for CHD-family remodelers in epigenetic silencing through a novel positive feedback mechanism where remodelers (Mit1) pair with chromatin modifying enzymes (Clr4) to promote H3K9me establishment and spreading.

## RESULTS

### Tethering Mit1 at an ectopic site induces transcriptional silencing

The canonical model for heterochromatin assembly in *S. pombe* involves sequence-specific factors that recruit Clr4 to establish H3K9me at pericentromeric repeats, telomeres, and the mating type locus. H3K9me subsequently serves as a binding platform to recruit diverse chromatin modifiers that repress transcription (Grewal, 2023). The CHD class remodeler, Mit1 is recruited to heterochromatin through its interaction with the HP1 protein Chp2, as part of a multi protein complex called SHREC (Job et al., 2016; Leopold et al., 2019b; Motamedi et al., 2008; Sugiyama et al., 2007). Since Mit1 is thought to function downstream of H3K9 methylation, we used a synthetic tethering approach to bypass its endogenous recruitment and potentially isolate the contribution of Mit1 from other factors involved in heterochromatin assembly (**Figure 1A**). Here, we fused Mit1 to the tetracycline repressor protein (*TetR-Mit1*) and targeted it to ten tetracycline operator sites (*10XtetO*) placed upstream of a reporter gene encoding NLS-GFP-mNeonGreen (*NLS-GFP-mNG*) (**Figure 1B**). We used fluorescence microscopy to assess whether cells displayed high (*GFP-ON*) or low (*GFP-OFF*) reporter gene expression. Qualitative analysis across three different biological replicates captured a subset of cells that exhibit a GFP-OFF state upon Mit1 tethering (**Figure 1C**). To quantify this effect, we performed flow cytometry and measured the fraction of GFP-OFF cells across three independent biological replicates. We observed substantial variability, with individual clones displaying ∼6%, 29%, and 20% GFP-OFF cells, respectively. In contrast, control experiments where we tethered an ATPase-dead Mit1 variant (*TetR-mit1^K587A^*) completely abolished silencing and all cells adopted a GFP-ON state (**Figure 1D**) (Creamer et al., 2014). To determine whether the GFP-OFF state established by Mit1 tethering is stable, we used a microfluidic device that allows for continuous imaging of individual cell lineages for extended periods of time (∼48 hours). Using this platform, we tracked individual cell lineages across ∼25 consecutive generations and confirmed that once established, the GFP-OFF state initiated by Mit1 tethering was continuously maintained across many generations (**Figure S1A**).

**Figure 1.**
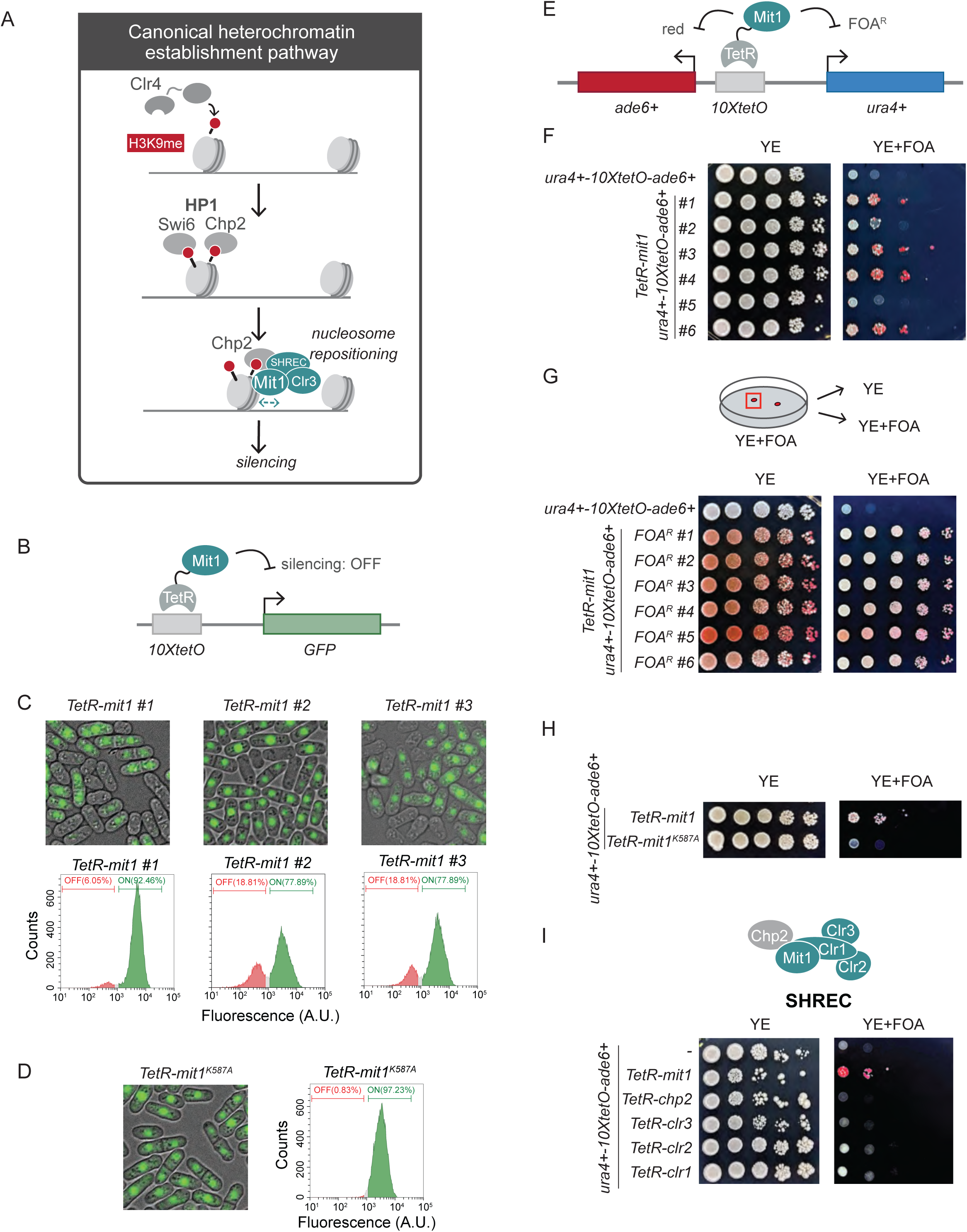
Tethering Mit1 at an ectopic site induces transcriptional silencing. (A) The canonical model for heterochromatin assembly in *S. pombe* involves sequence specific factors that recruit Clr4 to establish H3K9me. H3K9me serves as a binding platform for HP1 proteins (Swi6 and Chp2) to recruit chromatin modifiers that silence transcription. The CHD class remodeler, Mit1 is recruited to heterochromatin through its interaction with the HP1 protein Chp2, as part of a multi protein complex called SHREC (Clr1, Clr2, Clr3, Mit1 and Chp2). (B) Schematic of TetR-Mit1 recruitment to *10XtetO* binding sites placed upstream of a GFP reporter. (C) Fluorescence microscopy images of three independent *TetR-mit1* isolates in the *10XtetO-mNeonGreen-GFP* reporter strain (upper panel). Flow cytometry to quantify GFP expression (lower panel). Histograms quantifying GFP expression. GFP-OFF population (red); GFP-ON population (green) (D) Fluorescence microscopy images of *TetR-mit1^K587A^* expressing cells (left panel) in the *10XtetO-mNeonGreen-GFP* reporter strain. Flow cytometry to quantify GFP expression (right panel). Histogram quantifying GFP expression. GFP-OFF population (red); GFP-ON population (green). Mit1^K587A^ is an ATPase deficient mutant. (E) Schematic of TetR-Mit1 recruitment to *10XtetO* binding sites placed between *ade6+* and *ura4+* reporters (*ura4+ – ade6+* reporter). Silencing causes cells to turn red and enables cells to grow on FOA containing media (FOA^R^). (F) Silencing assay for six independent biological replicates of *TetR-mit1* in the *ura4+ – ade6+*reporter strain. (G) Silencing assay for the six TetR-Mit1 isolates from **1F** after FOA selection (FOA^R^) in the *ura4+ – ade6+* reporter strain. (H) Silencing assay of *TetR-mit1^K587A^* in the *ura4+ – ade6+* reporter strain. Mit1^K587A^ is an ATPase deficient mutant. (I) Silencing assay comparing TetR fused to each SHREC complex subunit (Clr1, Clr2, Clr3 and Mit1) in the *ura4+ – ade6+* reporter strain. For all plate-based assays, cells are plated at 10-fold serial dilutions on non-selective low adenine media (YE) and on low adenine media containing FOA (YE+FOA).

Because only a subset of cells established silencing upon Mit1 tethering, we developed a selection strategy to enrich for this population. We engineered a reporter system where *ura4+* and *ade6+* were inserted proximal to *10XtetO* binding sites (**Figure 1E**). When both reporters are silenced, colonies can grow on medium containing 5-fluoroorotic acid (FOA) (*ura4+* silencing) and accumulate a red pigment on plates containing low adenine (YE) (*ade6+* silencing). We plated equal numbers of cells on YE and YE+FOA plates. Since most cells fail to establish silencing*, ade6+* is expressed and colonies appear white when they grow on YE media (**Figure 1F**, left panel). Consistent with our GFP based assays, tethering TetR-Mit1 resulted in only a fraction of cells that were able to form red colonies which are resistant to FOA (∼1-10%) (**Figure 1F**, right panel). We also observed significant clone-to-clone variability wherein different numbers of colonies exhibit FOA resistance across independent biological replicates. Consistent with the stability of the GFP-OFF state based on our imaging experiments, re-plating FOA resistant colonies after selection led to the appearance of red colonies on both YE and YE+FOA plates across all replicates. Furthermore, all replicates exhibit increased survival on FOA medium suppressing all of the initial clone-to-clone variability (**Figure 1G**). Additionally, silencing required Mit1 catalytic activity, as the catalytic mutant failed to grow on YE+FOA media (**Figure 1H**, *TetR-mit1^K587A^*). These results suggest that silencing via Mit1 tethering occurs with variable efficiency, is dependent on its catalytic activity, and robust once cells overcome the initial establishment barrier.

Mit1 is a component of the SHREC complex, which combines both nucleosome remodeling and deacetylase functions (Sugiyama et al., 2007). To determine whether the silencing activity we observed is specific to Mit1, we individually tethered each subunit of the SHREC complex. Previous studies have shown that tethering the histone deacetylase, Clr3 leads to enhanced maintenance, but this process cannot initiate silencing (Raiymbek et al., 2020; Zofall et al., 2022). Indeed, tethering Clr3, Clr1, or Clr2 failed to induce silencing, and we observed no growth on YE+FOA plates (**Figure 1I**). Hence, the silencing establishment activity we observed was specific to tethering Mit1 and required its ATPase dependent remodeling function.

To ensure that silencing was not an artifact of FOA-based selection, we compared the frequency of red colonies on YE versus FOA-containing plates. We observed no difference in frequency between the two media types, indicating that silencing is not driven by FOA selection (**Figure S1B**). Furthermore, we observed the same level of stability for the OFF state regardless of whether colonies that establish silencing originate from YE or YE+FOA plates (**Figure S1C**). We also confirmed that silencing at the ectopic site depends exclusively on the tethered Mit1 allele which can fully complement endogenous Mit1 activity (**Figure S1D**).

Lastly, since Mit1 is a chromatin remodeler that repositions nucleosomes, we hypothesized that remodeler-dependent heterochromatin establishment would depend on nucleosome density. Consistent with this expectation, we observed that the deletion of Pob3 (part of the heterodimeric FACT complex) but not Mrc1, a chaperone involved replicative histone transfer, leads to a loss of silencing (**Figure S1E**) (Charlton et al., 2024; Holla et al., 2020; Yu et al., 2024).

### Mit1 tethering promotes H3K9 methylation establishment and spreading

We expected that TetR-Mit1 would remodel nucleosomes proximal to the tethering site and exert its influence primarily on reporter gene expression (GFP-mNeonGreen or *ura4+-ade6+*) without long-range effects (**Figure 2A**). However, RNA-sequencing revealed that loci extending well beyond the proximal *ura4+* and *ade6+* reporters are downregulated when Mit1 is tethered (**Figure 2B**). One possible mechanism in *S. pombe* that could contribute to the spread of silencing is Clr4-dependent H3K9 methylation (Ivanova et al., 1998; K. Zhang et al., 2008).

**Figure 2:**
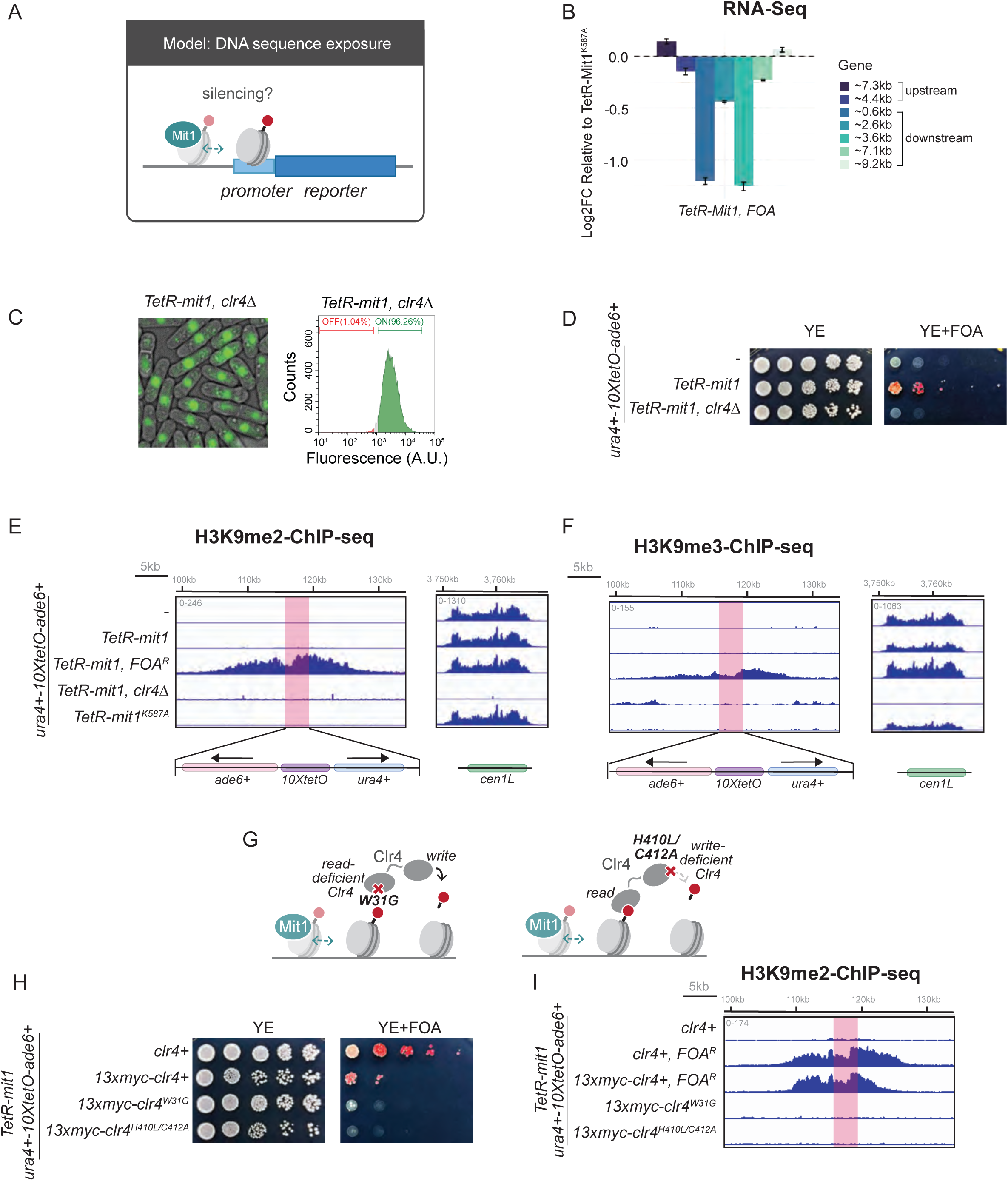
Mit1 tethering promotes H3K9 methylation establishment and spreading. (A) Schematic of how remodelers such as Mit1 reposition nucleosomes to regulate DNA sequences accessibility. (B) RNA-seq analysis of transcript levels in *TetR-mit1* FOA^R^ cells relative to the catalytic mutant (*TetR-mit1^K587A^*) across a 10kb region flanking the *10xTetO* binding site. Changes in expression are shown as log2 fold change normalized to *TetR-mit1^K587A^*(N=3). (C) Fluorescence microscopy images of *TetR-mit1, clr4Δ* (left panel) in the *10XtetO-mNeonGreen-GFP* reporter strain. Flow cytometry to quantify GFP expression (right panel). Histogram quantifying GFP expression. GFP-OFF population (red); GFP-ON population (green). (D) Silencing assay of TetR-Mit1, *clr4*Δ in the *ura4+ – ade6+* reporter strain. (**E,F**) ChIP-seq of H3K9me2 (**E**) and H3K9me3 (**F**) flanking the TetR-Mit1 binding site in the indicated genotypes. The *ura4+ – ade6* reporter which replaces the endogenous *ura4* locus is highlighted in red. A 35kb region spanning the tethering site is shown. (G) Schematic of Clr4 mutations leading to a defective chromodomain (*clr4^W31G^*) or a defective writing domain (*clr4^H410L/C412A^*). (H) Silencing assay measuring TetR-Mit1 silencing with Clr4 mutations *clr4^W31G^* or *clr4^H410L/C412A^* in the *ura4+ – ade6+* reporter strain background. (I) ChIP-seq of H3K9me2 surrounding the TetR-Mit1 binding site in the indicated genotypes. The *ura4+-10xtetO-ade6+* at the *ura4* locus is highlighted in red. A 35kb region surrounding the tethering site is shown.

Deletion of *clr4+* in cells with tethered Mit1 (*TetR-mit1, clr4Δ*) completely abolished silencing establishment, as evidenced by the loss of GFP silencing (all cells are GFP-ON) and the failure of cells to grow on YE+FOA media (**Figure 2C-D**). We performed chromatin immunoprecipitation followed by sequencing (ChIP-Seq) to map H3K9me2 and H3K9me3 levels in naïve cells where Mit1 is tethered, and FOA^R^ colonies (*TetR-mit1, FOA^R^*). While H3K9me2 and H3K9me3 levels are low and nearly undetectable in naïve cells, *TetR-mit1, FOA^R^* cells exhibit robust levels of H3K9me2 and H3K9me3 that spread across ∼20kb (**Figure 2E and Figure 2F**). In contrast, tethering the catalytically inactive Mit1 (*TetR-mit1^K587A^*) revealed no accumulation of H3K9me2 or H3K9me3, and it was not possible to pre-select colonies that are FOA resistant for subsequent assays.

The H3K9me domain size initiated by Mit1 rivals the extent of spreading that we typically observe upon directly tethering Clr4 (**Figure S2A**). Heterochromatin established by tethering Clr4 in *S. pombe* is robustly maintained in the absence of the heterochromatin antagonist, Epe1 (Audergon et al., 2015; Ragunathan et al., 2015). Cells can maintain silencing for ∼50-100 generations in an H3K9me-dependent manner without sequence-dependent establishment.

Similarly, *TetR-mit1, epe1Δ* cells exhibited a stronger maintenance phenotype (**Figure S2B**). Notably, we also observed weak maintenance in *epe1+* cells upon tethering Mit1 (*TetR-mit1, epe1+*), which could potentially represent a form of positional memory that depends on Mit1 remodeling activity similar to what has previously been noted in *S. cerevisiae* (Brothers and Rine, 2022). Taken together, these data suggest that tethering Mit1 promotes Clr4 recruitment and H3K9me spreading.

Next, we asked whether the read–write activity of Clr4 is necessary to reinforce and propagate Mit1-initiated silencing (K. Zhang et al., 2008). Clr4 has a chromodomain (CD) that binds to H3K9me nucleosomes and a SET domain that catalyzes H3K9me (**Figure 2G**) (Ivanova et al., 1998). Expressing the Clr4 chromodomain mutant (*clr4^W31G^*) or SET-domain mutant (*clr4^H410L/C412A^*) leads to the complete loss of silencing establishment (**Figure 2H**) (Iglesias et al., 2018; K. Zhang et al., 2008). Furthermore, ChIP-Seq measurements revealed no H3K9me accumulation in *TetR-mit1, clr4^W31G^* cells despite H3K9 methyltransferase activity being intact in this Clr4 mutant (**Figure 2I**). These data indicate that both H3K9me binding and catalysis is crucial for Mit1-initiated silencing unlike Clr4 tethering measurements where a CD deficient mutant can successfully establish functional heterochromatin (Audergon et al., 2015; Kagansky et al., 2009; Ragunathan et al., 2015).

### A systematic genetic screen identifies canonical heterochromatin factors required for Mit1-initiated silencing establishment

Because Mit1-initiated silencing depends on H3K9me, we performed a genome-wide screen to comprehensively define genetic dependencies associated with the silencing process. To accomplish this, we crossed strains containing *TetR-mit1* and the *ura4+ – ade6+* reporter to the *S. pombe* deletion library. The library consists of 2,988 non-essential gene deletions marked with *kanMX6* but excludes most genes encoding mitochondrial proteins as well as deletions that cause severe growth defects. After crossing and selection, cells were plated on media containing FOA to quantitatively measure how each deletion affects silencing establishment (**Figure 3A**). We compared our TetR-Mit1 screen results to a previous genetic screen that was carried out using a *ura4+* reporter at the silent mating type (*mat)* locus (Muhammad et al., 2024). This comparison identified distinct clusters of genes that influence Mit1-initiated silencing. The largest and most informative cluster (Cluster I) consists of factors required for both Mit1-dependent silencing and silencing at the mating type locus capturing substantial overlap between ectopic Mit1 targeting and endogenous heterochromatin pathways (**Figure 3B**). Strikingly, these genes belong to the heterochromatin core machinery with well-characterized roles in silencing. The remaining clusters (Clusters II-VII) captured several differences between the ectopic and endogenous contexts, but these findings did not affect our primary assertion that tethering Mit1 leads to H3K9 methylation dependent heterochromatin establishment (**Figure S3A-G**).

**Figure 3.**
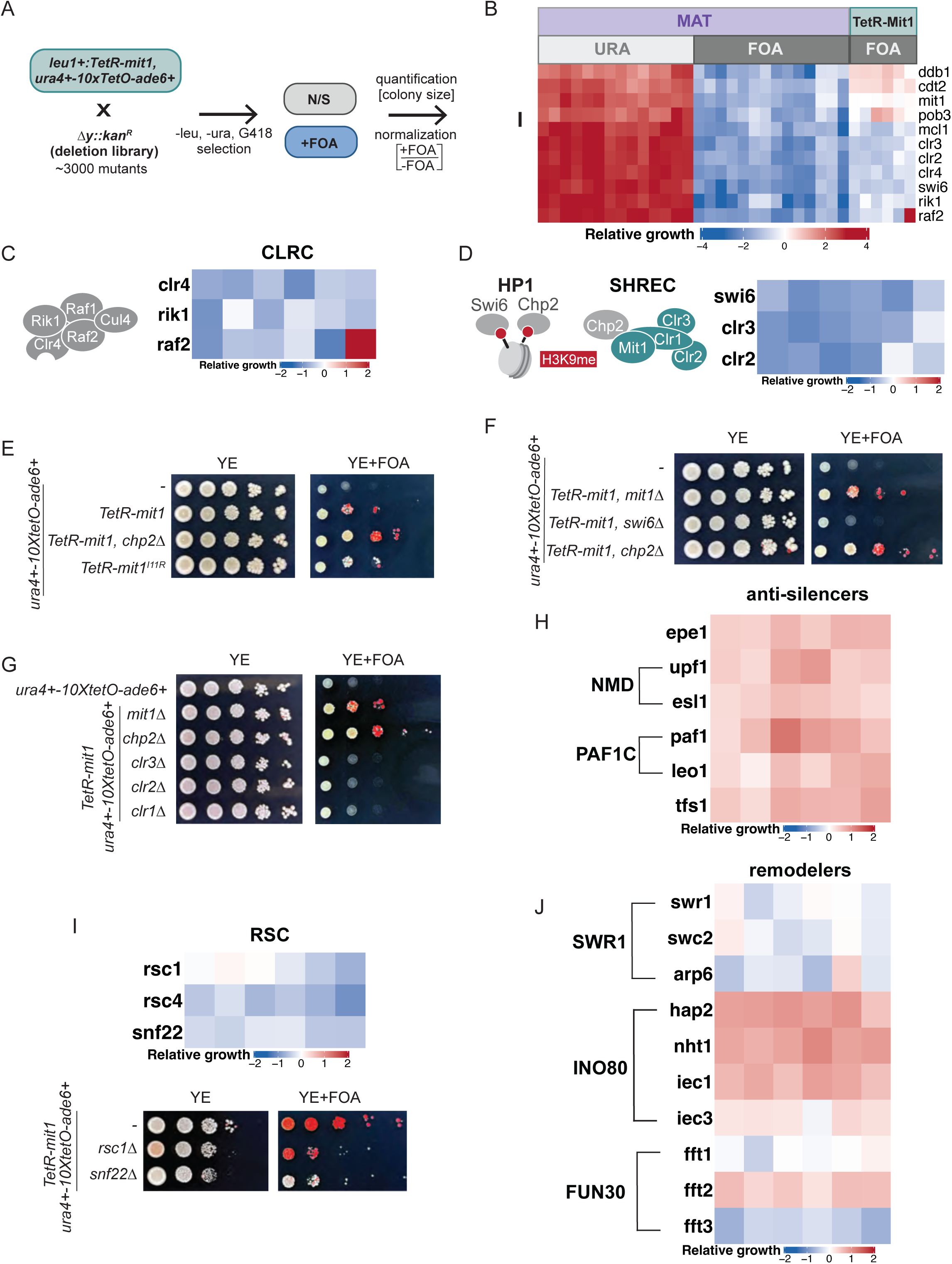
A systematic genetic screen identifies canonical heterochromatin factors required for Mit1-initiated silencing establishment. (A) Schematic of screening strategy to cross *TetR-mit1, ura4+ – ade6+* reporter with the yeast deletion library and perform silencing based selection assays. (B) Heatmap (Cluster-I) displaying comparisons of reporter activity between core-heterochromatin protein deletions in a *ura4+* reporter at the endogenous heterochromatin *mat* locus and in the *ade6+-ura4+* reporter in *TetR-mit1* cells. (C) Heatmap displaying relative growth scores for CLRC complex subunits. (D) Heatmap displaying relative growth scores for HP1 and SHREC complex subunits. (E) Silencing assay measuring TetR-Mit1 silencing in *chp2Δ* cells or in cells where the Chp2-Mit1 interaction is disrupted (*TetR-mit1^I11R^*). (F) Silencing assay measuring TetR-Mit1 silencing in the absence of HP1 proteins (*swi6Δ* versus *chp2Δ*). (G) Silencing assay measuring TetR-Mit1 silencing in the absence of SHREC proteins (*clr1Δ, clr2Δ, clr3Δ* and *chp2Δ)*. (H) Heatmap displaying relative growth scores for anti-silencing factors in *TetR-mit1* strains. (I) Heatmap displaying relative growth scores for RSC in *TetR-mit1* strains (top panel). Silencing assay measuring silencing in the absence of antagonizing chromatin remodelers, Rsc1 and Snf22 (bottom panel). (J) Heatmap displaying relative growth scores for remodeler complexes in *TetR-mit1* strains. Relative growth values represent individual colony sizes of mutants on uracil-lacking and FOA-containing media normalized to non-selective growth conditions and are log2-transformed. The results represent six replicates from three independent genetic screens with each with two technical replicates. All silencing assays were performed using the *ura4+ – ade6+* reporter strain.

Consistent with the *clr4Δ* phenotype, multiple components of the CLRC complex (Clr4, Rik1, Raf2) were found to be essential for Mit1-initiated silencing (**Figure 3C**). The other two members of CLRC, Raf1 and Cul4, were not included in the screen. The deletion of other canonical heterochromatin factors also exhibited a loss of silencing such as the HP1 protein, Swi6, and members of the SHREC complex, Clr2 and Clr3 (**Figure 3D**) (Motamedi et al., 2008; Sadaie et al., 2008; Yamada et al., 2005). The second HP1 protein Chp2 and a structural component of SHREC, Clr1, were not included in the screen.

We validated the genetic screen with targeted deletions and mutations of individual factors (including those that are missing in the screen). Disrupting the Chp2-Mit1 interaction– either by deleting Chp2 (*chp2Δ*) or by expressing a Mit1 point mutant (*TetR-mit1^I11R^*) that abolishes their interaction had no effect on silencing establishment, consistent with our expectation that tethering Mit1 bypasses endogenous recruitment mechanisms (**Figure 3E**)(Leopold et al., 2019a). In contrast, the targeted deletion of the other HP1 protein Swi6 and all components of the SHREC complex (Clr1, Clr2, or Clr3) led to the complete loss of silencing establishment upon tethering Mit1 (**Figure 3F and 3G**) (Sugiyama et al., 2007). Hence, Mit1-initiated silencing recapitulates the same genetic dependencies as canonical or endogenous heterochromatin domains, with the notable exception that Chp2 is dispensable when Mit1 is tethered.

The screen also identified several anti-silencing factors that antagonize canonical heterochromatin such as the H3K9 demethylase, Epe1, members of the NMD pathway, Upf1 and Esl1, and factors involved in RNA polymerase II elongation (Paf1C) (**Figure 3H**) (Ayoub et al., 2003; Bhatt et al., 2026a; Kowalik et al., 2015; Verrier et al., 2015; Zofall and Grewal, 2006). Deleting subunits involved in anchoring the histone acetyltransferase complex, Mst2C to H3K36me3 lead to enhanced silencing. Individual subunits of the other major acetyltransferase complex, SAGA produce distinct effects with some subunits promoting silencing while others weaken this process (**Figure S3H and S3I**). This observation suggests distinct functions of submodules within the complex in line with previous reports (Bao et al., 2019; Flury et al., 2017; Georgescu et al., 2020; Reddy et al., 2011).

One possibility is that Mit1-initiated silencing is simply an outcome of antagonism with opposing remodelers such as RSC-family complexes which are specifically excluded from deacetylated heterochromatin domains (Sahu et al., 2024). Deleting chromatin remodelers, Snf22 and Rsc1, failed to enhance silencing but rather produced mild silencing defects in the screen (**Figure 3I**). The deletion of other remodeler complexes such as SWR1 had no effect on silencing, INO80 deletions enhanced silencing as previously noted, and the FUN30 remodeler deletions produced disparate effects suggesting that these paralogs have distinct functions (**Figure 3J**) (Hou et al., 2010; Shan et al., 2020; Taneja et al., 2017).

### Clr4 dosage determines the frequency of Mit1-initiated heterochromatin establishment

Our screen revealed that the deletion of factors involved in sequence specific heterochromatin establishment at the pericentromeres and the subtelomeres improved Mit1-initiated silencing (**Figure 4A**) (Tadeo et al., 2013). These included factors involved in the RNAi pathway such as, Tas3 and Chp1, which are part of the RITS complex, and Ers1 and Cid12, which are part of the RDRP complex (Motamedi et al., 2004; Verdel et al., 2004). In addition, we identified telomere-specific binding proteins such as Tlh2, the shelterin subunits Rap1, Poz1, and Ccq1 (**Figure 4B-C**) (Kanoh et al., 2005; Wang et al., 2016). We validated these findings with the targeted deletion of a shelterin subunit, Ccq1 (*ccq1Δ*) (**Figure S4A**). We confirmed that the enhanced silencing was due to an increase in H3K9me2 and H3K9me3 at the reporter locus in *ccq1Δ* cells without the requirement for prior FOA selection (**Figure S4B and S4C**).

**Figure 4.**
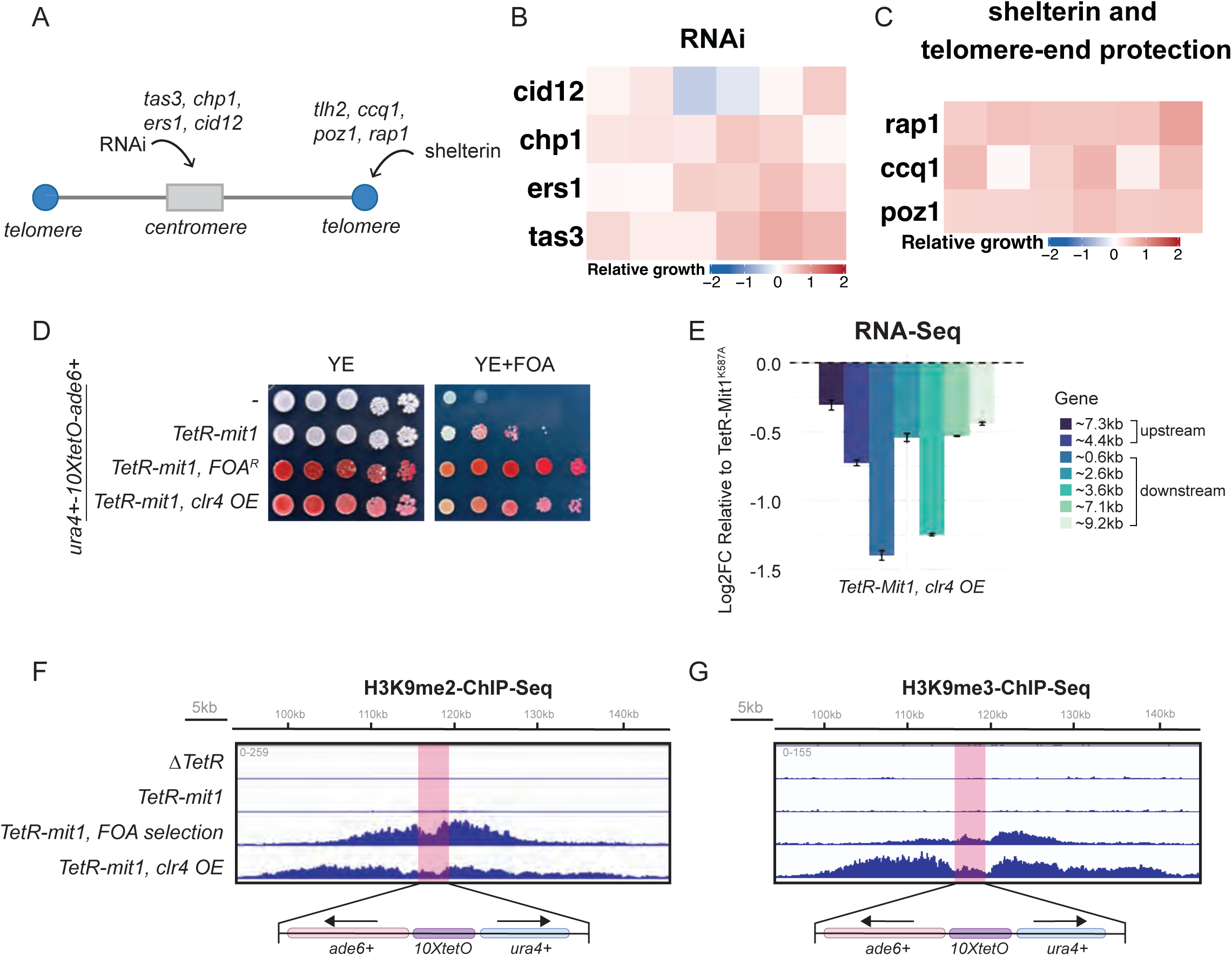
Clr4 dosage determines the frequency of Mit1-initiated heterochromatin establishment. (A) Schematic of RNAi and associated factors acting at centromeres and shelterin and associated factors acting at telomeres. (B) Heatmap displaying relative growth scores for deletions of factors involved in the RNAi pathway. (C) Heatmap displaying relative growth scores for deletions of shelterin subunits involved in establishing heterochromatin at telomeres and telomere-end protection. (D) Silencing assay measuring TetR-Mit1 silencing in Clr4 overexpression. (E) RNA-seq analysis of transcript levels in *TetR-mit1 clr4 OE* cells relative to the catalytic mutant (*TetR-mit1^K587A^*) across a 10kb region flanking the *10xTetO* binding site. Changes in expression are shown as log2 fold change normalized to *TetR-mit1^K587A^*(N=3). (**F, G**) ChIP-seq of H3K9me2 (**F**) and H3K9me3 (**G**) flanking the TetR-Mit1 binding site in the indicated genotypes. The *ura4+ – ade6* reporter which replaces the endogenous *ura4* locus is highlighted in red. A 50kb region spanning the tethering site is shown. All silencing assays were performed using the ura4+ – ade6+ reporter strain.

We hypothesized that endogenous heterochromatin loci sequester Clr4, which makes H3K9me deposition the rate-limiting step for Mit1 to initiate silencing. To test this hypothesis, we overexpressed *clr4* using the *adh15* promoter, which leads to a ∼4-fold increase in RNA levels compared to endogenous *clr4* (*clr4 OE*) (**Figure S4C**). Strikingly, *clr4 OE* led to most cells establishing heterochromatin, which can be readily visualized as colonies that appear red on both YE and YE+FOA plates (left panel, **Figure 4D**). The fraction of cells that exhibit FOA resistance was significantly higher in cells where Clr4 was overexpressed. RNA-seq revealed a more extensive domain of silencing and ChIP-Seq of H3K9me2 and H3K9me3 exhibited substantially increased spreading (∼40kb) without any need for FOA selection (**Figure 4E-G**). These findings support the idea that the dosage of heterochromatin factors broadly and Clr4 specifically tunes the silencing establishment frequency upon Mit1 tethering.

### Mit1 overexpression promotes ectopic heterochromatin establishment

Having established that Mit1 remodeling activity is sufficient to promote Clr4 recruitment and H3K9 methylation, we next asked how this activity alters heterochromatin establishment at endogenous loci. We replaced the endogenous *mit1* promoter with a moderate overexpression promoter (*adh11*) and assessed its effect on heterochromatin spreading using an *ade6+* reporter positioned just outside the mating type boundary sequences (**Figure 5A**, *mat2P::ade6+*) (Iglesias et al., 2018; Zofall et al., 2016). Deleting the H3K9me eraser Epe1 (*epe1Δ*) in this reporter background leads to increased H3K9me spreading across boundary sequences and produces uniformly red colonies. In contrast, *epe1Δ, mit1 OE* cells exhibit an unstable silencing phenotype. In this case, we observed both red and white colonies. We also noted substantial clone-to-clone variability in silencing phenotypes between different isolates (**Figure 5B**).

**Figure 5.**
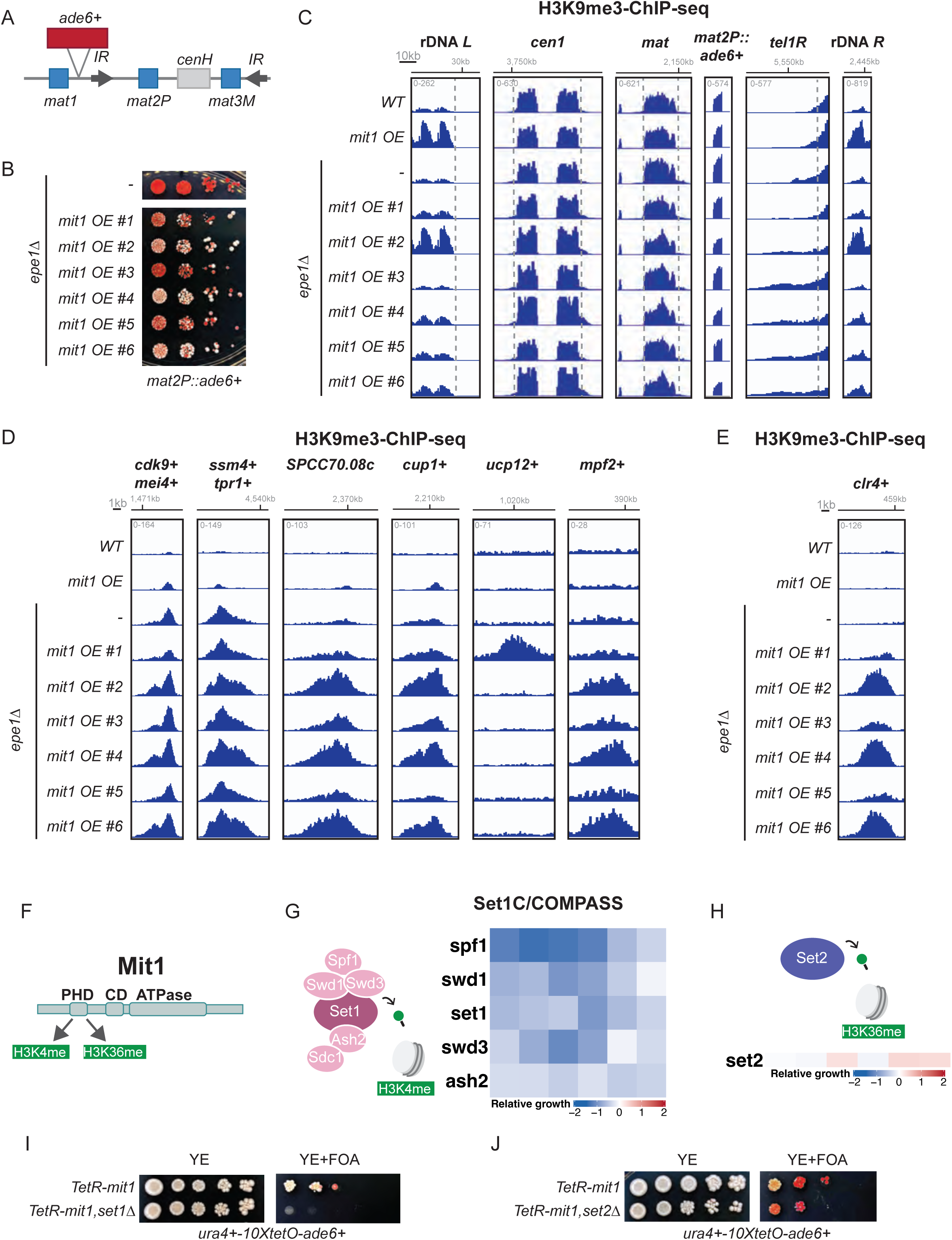
Mit1 overexpression promotes ectopic heterochromatin establishment. (A) Schematic of the mat locus with an ade6+ reporter near the mating type locus (*mat2P::ade6+*). (B) Silencing assay of *mat2P::ade6+* reporter strains in the indicated genotypes. (C) ChIP-seq of H3K9me3 at endogenous heterochromatin regions (in the indicated genotypes. (D) ChIP-seq of H3K9me3 at regions with *de novo* peaks in Mit1 OE. (E) ChIP-seq of H3K9me3 at the *clr4+* locus. (F) Schematic of domains in Mit1: the PHD domain binds to H3K4me3 and H3K36me3 *in vitro*, the CD domain binds DNA, and the C-terminus harbors the catalytic ATPase domain. (G) Heatmap displaying relative growth scores for SET1C/COMPASS deletions in *TetR-mit1* expressing cells in the *ura4+ – ade6+* reporter strain background. (H) Heatmap displaying relative growth scores for Set2 deletion in *TetR-mit1* expressing cells in the *ura4+ – ade6+* reporter strain background. (I) Silencing assay of *TetR-mit1, set1Δ* in the *ura4+ – ade6+* reporter strain background. (J) Silencing assay of *TetR-mit1, set2Δ* in the *ura4+ – ade6+* reporter strain background.

To determine the molecular basis for the observed clone-to-clone variability, we performed ChIP-seq to map H3K9me2 and H3K9me3 genome-wide. At constitutive heterochromatin sites, Mit1 overexpression had region-specific effects. This includes a substantial increase in H3K9me3 at the rDNA locus upon Mit1 overexpression, which exhibits no additional increase in *epe1Δ, mit1OE* cells. We observed a modest increase of H3K9me3 at the pericentromeres, but substantial spreading of H3K9me3 at the telomeres across all replicates (**Figure 5C and S5A**). The increase in spreading is additive in an *epe1Δ* background, suggesting that Mit1 facilitates H3K9me deposition through a mechanism that is distinct from Epe1. At facultative heterochromatin sites — including meiotic genes such as *mei4+* and *ssm4+* and heterochromatin islands at *SPCC70.08c*, *cup1*, *ucp12*, and *mpf2* — we detected H3K9me3 peaks that substantially exceeded the levels observed in *epe1Δ* cells alone (**Figure 5D and S5B**)(Torres-Garcia et al., 2020; Zofall et al., 2012).

Most strikingly, we observed accumulation of both H3K9me2 and H3K9me3 at the *clr4+* locus itself in all *epe1Δ, mit1 OE* clones (**Figure 5E and S5C**). The level of H3K9me at the *clr4+* locus qualitatively maps to the degree of variegation in silencing (white versus red colonies) of the *mat2P::ade6+* reporter, with clones that have more white colonies exhibiting higher levels of H3K9me at *clr4+*. This is consistent with previous work showing that heterochromatin misregulation can trigger adaptation, wherein cells silence *clr4+* to mitigate uncontrolled H3K9me spreading (Wang et al., 2015). Notably, *epe1Δ* strains alone showed no H3K9me accumulation at *clr4+*, indicating that this adaptive silencing process is additive only in a Mit1 overexpression context.

We next considered how Mit1 is recruited to genomic regions outside of heterochromatin suggesting that there might be a Chp2-independent binding mechanism that regulates Mit1 activity. Mit1 has a PHD domain that *in vitro* binds H3K4me3 and H3K36me3 peptides (**Figure 5F**)(Creamer et al., 2014). Our genetic deletion screen highlighted that all subunits of the Set1C complex that are present in the library and are critical for H3K4me3 deposition (Set1, Ash2, Spf1, Swd1, Swd3), are necessary for Mit1-initiated silencing (**Figure 5G**) (Mikheyeva et al., 2014; Noma and Grewal, 2002). Deleting Set2 which deposits H3K36me3 had no effect on silencing suggesting that H3K36 methylation is dispensable for this process (**Figure 5H**) (Georgescu et al., 2020). These findings were confirmed by targeted deletions of Set1 and Set2, which corroborated our screen results (**Figure 5I and 5J**). These data suggest that H3K4me regulates Mit1 activity, and this regulatory function could serve as an anchor for its recruitment to novel sites in the genome to induce *de novo* heterochromatin establishment.

### Mit1 promotes H3K9me establishment through remodeling activity

We envisioned two possibilities for how Mit1 promotes H3K9me establishment: **1)** directly through physical interactions involving the CLRC complex (**Figure 6A**) (Hong et al., 2005; Horn et al., 2005) or **2)** indirectly through chromatin remodeler activity that promotes Clr4 recruitment (**Figure 6B**). To test the direct interaction model, we used existing structural data to truncate Mit1 at its N-terminus (eliminates Chp2 interaction site) (Leopold et al., 2019a) or at its C-terminus (eliminates SHREC interaction site via Clr1) (Job et al., 2016). While the N-terminus is dispensable for silencing establishment, the C-terminus is required, indicating that Mit1 functions in coordination with other SHREC complex subunits to promote heterochromatin establishment (**Figure 6C**). These results are also consistent with every subunit of SHREC being essential for Mit1-initiated silencing (**Figure 1I**).

**Figure 6.**
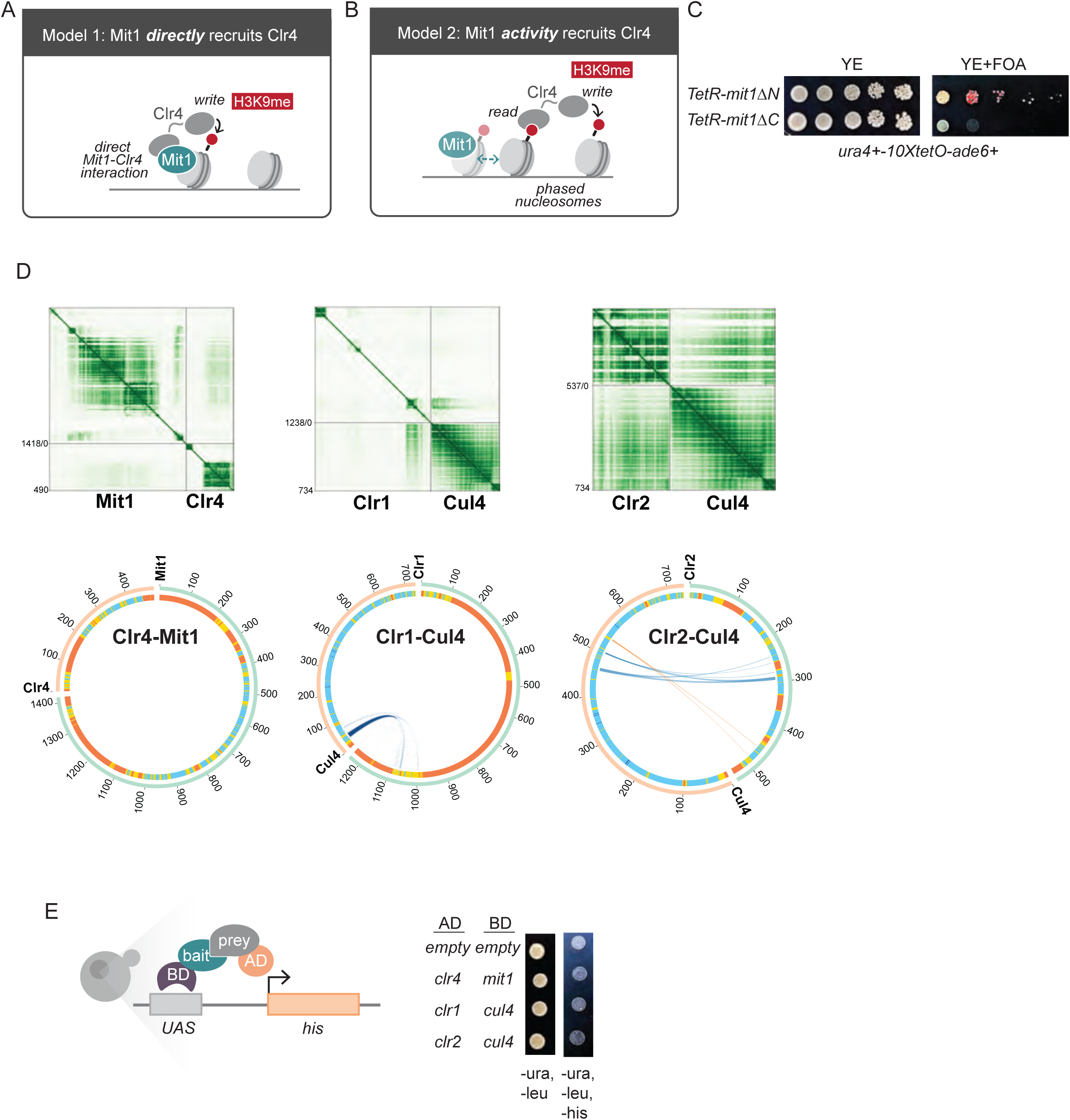
Mit1 facilitates Clr4 recruitment through an indirect mechanism. (A) Schematic of model describing direct interaction between Mit1 and Clr4. (B) Schematic of model describing indirect interaction between Mit1 and Clr4. Mit1 spaces nucleosomes in a specific configuration that promotes Clr4 recruitment. (C) Silencing assay of TetR-mit1 with C or N terminus truncations in the *ura4+ – ade6+* reporter strain background. (D) AlphaBridge depicting interacting residues of the two pairs predicted to have interactions in addition to Clr4-Mit1. (E) Yeast two-hybrid (Y2H) to test interaction between predicted SHREC-CLRC interaction pairs. Figure 7: Mit1 remodeling activity promotes Clr4 activity.

Next, we used AlphaFold server (based on AlphaFold3) to screen *in silico* for potential pair-wise interactions between four members of SHREC (Clr1, Clr2, Clr3, and Mit1) and all the five members of CLRC (Clr4, Raf1, Raf2, Rik1 and Cul4) (Abramson et al., 2024). The AlphaFold3 predictions were evaluated using AlphaBridge which enabled us to identify and visualize interactors based on default confidence metrics (Álvarez-Salmoral et al., 2025). We noted two predicted interactions, Clr1-Cul4 and Clr2-Cul4 (**Figure 6D**). To validate the AlphaFold3 predictions, we performed yeast two-hybrid assays by co-expressing these protein pairs as well as Clr4-Mit1 in budding yeast (Fields and Song, 1989). We observed no growth on selective medium with any of these potential interactor pairs suggesting that these were likely false positive predictions (**Figure 6E,** *SC-ura-leu-his*). These findings are also fully consistent with earlier work in which IP-MS of individual SHREC complex subunits fails to detect an interaction with CLRC, suggesting that these complexes function independently to regulate H3K9 methylation (Hong et al., 2005; Horn et al., 2005; Motamedi et al., 2008; Sugiyama et al., 2007). While we cannot exclude the possibility of transient or context-dependent interactions, the absence of evidence based on our structural and genetic data argues against a stable physical connection between SHREC (Mit1) and CLRC (Clr4). Therefore, we favor a model in which Mit1 remodeling promotes H3K9me establishment through a chromatin based mechanism, by positioning nucleosomes in a configuration that favors Clr4 read-write function.

## DISCUSSION

Our current understanding of chromatin remodelers is premised on the notion that these enzymes primarily function by repositioning nucleosomes to regulate access to the underlying DNA sequence (Tsukiyama, 2002). Here, we demonstrate that the CHD-family remodeler Mit1 can establish a repressive chromatin state by nucleating broad domains of H3K9me (∼20kb), which are dependent on Clr4 and the ATPase activity of Mit1. Several findings are consistent with our chromatin-based model of Clr4 recruitment. First, the catalytic mutant of Mit1 fails to establish silencing or H3K9me when tethered (**Figure 1D and 1H**), arguing against a purely structural role for Mit1 in this process. Second, the chromodomain mutant of Clr4 is unable to establish H3K9me in cells where Mit1 is tethered despite having intact catalytic activity, yet directly tethering the same chromodomain mutant produces uniformly silenced colonies with high levels of H3K9me (Audergon et al., 2015; Kagansky et al., 2009; Ragunathan et al., 2015). Third, we observe highly variable silencing establishment frequencies (**Figure 1**) suggesting that a stochastic or transient H3K9me event could be the rate limiting step during nucleation.

This distinction argues that Mit1 does not recruit Clr4 through a direct physical interaction but instead suggests that the nucleosome remodeling activity of Mit1 amplifies heterochromatin establishment by a chromatin-based mechanism that stimulates Clr4 recruitment.

Tethering Mit1 uncovers mechanistic insights that are otherwise challenging to parse at endogenous heterochromatin loci. Rather than acting as the final step in silencing, we propose that Mit1 is integral to the read-write positive feedback loop: nucleosome remodeling by Mit1 promotes Clr4 read-write activity, which in turn leads to H3K9me deposition, recruiting HP1 proteins such as Chp2, which in turn recruit additional copies of Mit1 (**Figure 7**). Because this loop is self-reinforcing, increasing Mit1 dosage leads to enhanced H3K9me3 spreading at telomeres beyond what is observed in *epe1Δ* cells alone (**Figure 5C**). However, Mit1 overexpression is clearly detrimental to heterochromatin stability since it provokes an adaptive epigenetic response that most prominently leads to *clr4+* silencing (Wang et al., 2015).

**Figure 7.**
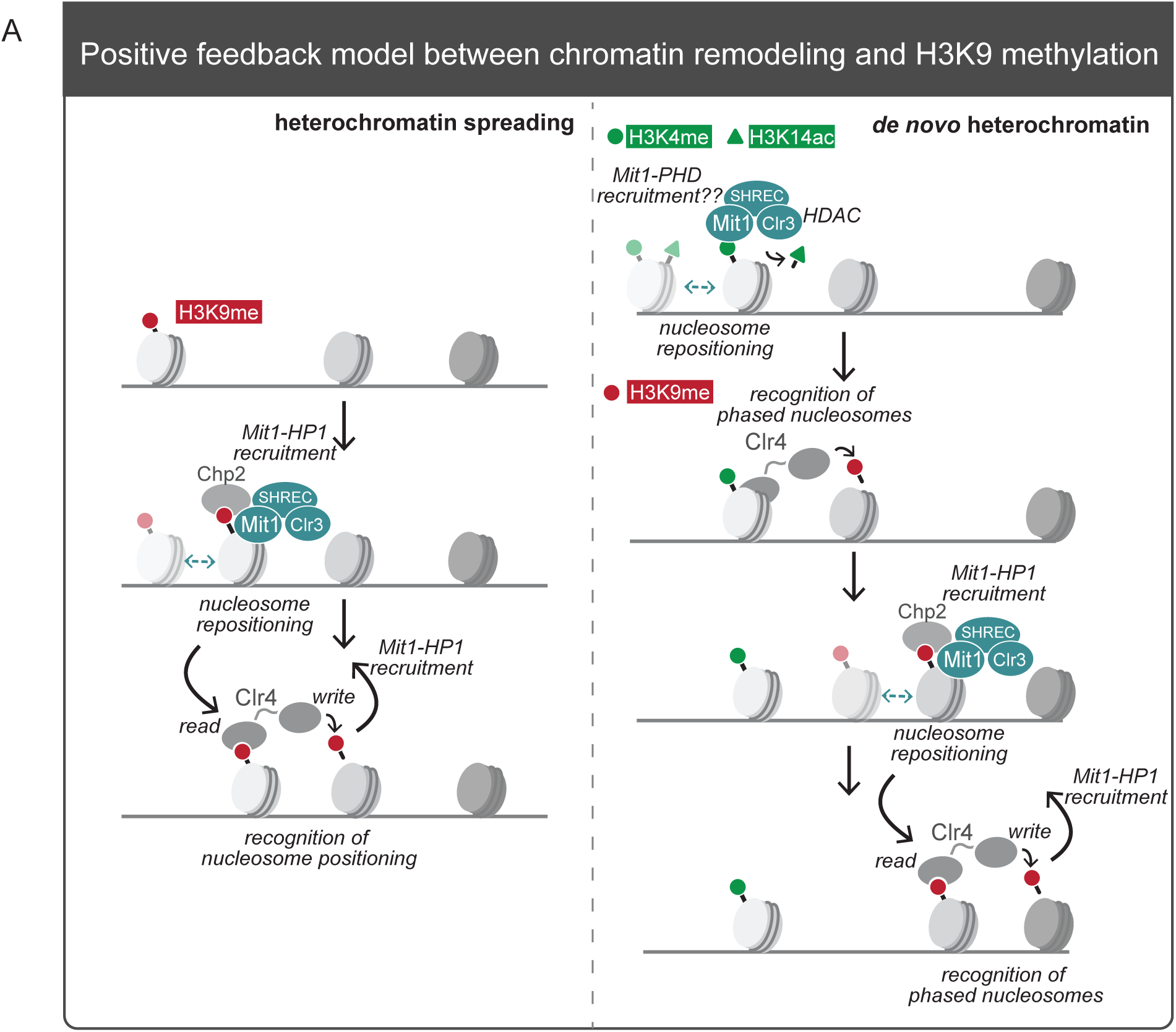
Model: Mit1 is part of a positive feedback loop where changes in nucleosome organization influence H3K9 methylation establishment and spreading. (A) *Heterochromatin spreading*: At endogenous heterochromatin, Mit1 is recruited to H3K9 methylated nucleosomes via its interaction with Chp2. Mit1 then repositions nucleosomes in a configuration that allows for Clr4 read-write activity. After H3K9me deposition, Chp2 binds and recursively recruits Mit1. This allows for more efficient spreading of H3K9me consistent with the notion that methyltransferase activity is sensitive to di-nuclosome spacing and orientation. (B) *De novo heterochromatin*: Mit1 is recruited to euchromatin, potentially through interactions involving its PHD domain binding to H3K4 methylation. This binding likely regulates its activity and recruitment. Mit1 then repositions nucleosomes in a configuration that promotes Clr4 read-write activity. This in turn recruits Chp2 and additional Mit1 across a domain. This allows for *de novo* heterochromatin formation.

Nucleosome spacing and geometry are critical determinants of methyltransferase activity as reflected in structural studies of PRC2 in complex with di-nucleosome substrates (Poepsel et al., 2018). Recent structural studies also show that Clr4 adopts a similar configuration where the enzyme bridges a di-nucleosome substrates, with its chromodomain binding to an H3K9-methylated nucleosome and its SET domain binding to and methylating adjacent unmodified nucleosomes (Saab et al., 2024). Based on these mechanistic observations, we envision that Mit1 promotes H3K9 methylation by remodeling chromatin to generate di-nucleosome configurations with the right spacing and orientation that enhance Clr4 activity. In contrast to chromatin remodeler complexes, such as SWI/SNF, RSC, or BAF that evict or dismantle nucleosomes to reset epigenetic states, Mit1 and by extension other CHD remodelers likely reposition nucleosomes to adopt configurations that can be exploited by diverse chromatin modifying enzymes (Hodges et al., 2016). In the case of *S. pombe,* we propose that Clr4 exploits the nucleosome spacing that Mit1 sets up to establish and spread H3K9 methylation.

While Mit1 catalytic activity is necessary to initiate silencing in our ectopic system, this process requires all components of the SHREC complex. SHREC contains the histone deacetylase Clr3, and tethering-induced silencing is abolished in *clr3Δ* cells (**Figure 3H**). The most parsimonious explanation for our data is that Mit1-initiated silencing depends on both nucleosome remodeling and histone hypoacetylation (Tong et al., 1998). Although structural data suggests only loose coupling between both activities, our genetic data strongly supports coordination since the C-terminus of Mit1 is essential to establish silencing (**Figure 6B**) (Job et al., 2016). This logic of remodelers stimulating histone methyltransferase activity may be broadly conserved as previously observed in the case of NuRD, the remodeler-deacetylation complex in humans which appears to recruit PRC2 to a subset of target genes in embryonic stem cells (Reynolds et al., 2012). In essence, our studies lend strong support to a model where NuRD enzymatic activity rather than sequence-specific recruitment may serve as the primary signal for PRC2 recruitment in some genomic contexts although this finding remains to be formally tested.

Beyond its role at endogenous sites, our data reveal that Mit1 can promote *de novo* H3K9me establishment in a sensitized background. Strikingly, these *de novo* H3K9me peaks occur at *clr4+* and *cup1+* which are loci linked to adaptive responses during heterochromatin misregulation or caffeine resistance (Torres-Garcia et al., 2020; Wang et al., 2015). Recent studies identified a heterochromatin regulatory hub (HRH) in which Raf1 levels, modulated by environmentally responsive pathways, are rate-limiting for CLRC assembly (Bhatt et al., 2026b). The dosage sensitivity of Mit1-initiated silencing in our studies raises the possibility that *de novo* establishment by Mit1 can be triggered when CLRC levels increase under stress conditions.

Together, our results suggest a two-step model for adaptive heterochromatin formation: Mit1 initiates *de novo* H3K9me domains by repositioning nucleosomes which in turn can be exploited by Clr4 and the Raf1-dependent HRH pathway to establish H3K9me that is stably propagated or allowed to decay depending on the environment. This architecture couples remodeler-driven initiation to an environmentally regulated maintenance switch, linking nucleosome organization to adaptive epigenetic responses. Our results do not negate the importance of sequence-specific establishment factors at endogenous heterochromatin loci. Instead, we argue that remodelers provide a unique source of positive feedback that amplifies Clr4 read-write activity which might play a role in sequence-independent heterochromatin establishment during stress.

A central tenet of epigenetics is that readers and writers function as coordinated pairs as part of a self-reinforcing mechanism that enables histone modifications to be propagated across multiple generations (Allshire and Madhani, 2018). Our findings identify a distinct but complementary partnership between a chromatin remodeler and a writer, suggesting that remodeler–writer pairs may represent a new frontier for understanding chromatin regulation.

One important implication of our findings is that remodelers are not inherently activating or silencing. Instead, remodeler activity can position nucleosomes in configurations that can be exploited by different chromatin modifying enzymes or reader proteins. In *S. pombe,* tethering Mit1 leads to repositioned nucleosomes that adopt a configuration that facilitates Clr4 mediated H3K9me establishment. In plants, the CHD-family remodeler PICKLE is required for PRC2 mediated H3K27me spreading, and in mammals, the NuRD complex similarly promotes PRC2 recruitment (Liang et al., 2024; Reynolds et al., 2012). These observations suggest that the framework of writer-remodeler pairs may define a conserved regulatory logic that facilitates the establishment and maintenance of epigenetic states across eukaryotes. We anticipate that tethering remodelers in mammalian cells will also lead to the establishment of epigenetic silencing by nucleating novel histone modification states *de novo* (Ornelas et al., 2026). If this logic extends to mammalian systems, remodeler-initiated silencing could contribute to the acquisition of chemotherapeutic resistance by silencing novel gene targets by initiating heterochromatin establishment in a DNA sequence independent manner (Flavahan et al., 2017; Guler et al., 2017; Sharma et al., 2010) .

## ACKNOWLEDGMENTS

We thank members of the Ragunathan lab for their insightful feedback in helping edit the manuscript and interpret the data: Neha Arora, Emily Metcalf, Katelyn Strauss, and Elizabeth Hemenway. We thank Danesh Moazed for sharing fission yeast strains used in this study. This work was supported by an NIH/NIGMS Award (R35GM137832) to K.R., an American Cancer Society Award (RSG-22-117-01-DMC) to K.R., and the Heisenberg Programme by the German Research Foundation (DFG, project ID 464293512) to S.B.

## DECLARATION OF GENERATIVE AI USAGE

Generative artificial intelligence (AI) tools (ChatGPT, OpenAI) were used solely for language editing and text refinement during manuscript preparation. All scientific content, data interpretation, and conclusions were generated, reviewed, and verified by the authors, who take full responsibility for the accuracy and integrity of the work.

## RESOURCE AVALIABILITY

Further information and requests for resources and reagents should be directed to and will be fulfilled by the lead contact, Kaushik Ragunathan

### Materials availability

All unique/stable reagents generated in this study are available from the lead contact without restriction.

### Data and code availability

ChIP-seq and RNA-seq data have been deposited at GEO and are publicly available as of the date of publication. ChIP-Seq accession number is GSE325535. RNA-Seq accession number is GSE325547. This paper does not report original code. Any additional information required to reanalyze the data reported in this paper is available from the lead contact upon request.

## AUTHOR CONTRIBUTIONS

M.S. conceptualized the project, developed the methodology, performed the investigation, analyzed the data, visualized the data, and reviewed/edited the final manuscript, A.C.L. developed the methodology, performed the investigation, analyzed the data, visualized the data, and reviewed/edited the final manuscript, A.M. performed the investigation, analyzed the data, and reviewed/edited the final manuscript, L.W. performed the investigation and analyzed the data, and reviewed/edited the final manuscript, F.H. conceptualized the project, developed the methodology, A.Z.A developed the methodology, visualized the data, and reviewed/edited the final manuscript, S.B. conceptualized the project, analyzed the data, visualized the data, reviewed/edited the final manuscript, and supervised the study, K.R. conceptualized the project, analyzed the data, visualized the data, wrote the original draft, reviewed/edited the final manuscript, and supervised the study,

## METHODS

### Plasmid

Plasmids containing *10XtetO* binding sites upstream *ade6+* and downstream *ura4+* were constructed by using Gibson to insert *ura4+* into a *10xtetO-ade6+* plasmid using inverse PCR. The plasmid containing *adh11-2xFLAG-TetR-mit1* was constructed by amplifying *adh11* and *2xFLAG-TetR* from plasmid DNA and amplifying *mit1* from genomic DNA with overhangs and assembled into a pDual plasmid using Gibson. Subsequently *pDual-adh11-2xFLAG-mit1* (Mit1 OE plasmid), *pDual-adh11-TetR-2xFLAG-mit1^K587A^, pDual-adh11-TetR-2xFLAG-mit1^ΔC^, pDual-adh11-TetR-2xFLAG-mit1^ΔN^,* were made by doing site-directed mutagenesis and inverse PCR on the *pDual-adh11-TetR-mit1* plasmid. All plasmids were confirmed using whole plasmid sequencing.

### Strains

The strain containing *ade6+-10xTetO-ura4+* was made using sequences amplified from *S.pombe* genomic DNA. Cells with a *ura4-D18* allele were then transformed using a standard electroporation transformation approach and selected with FOA. Subsequently, to insert *adh11-TetR-mit1,* the pDual plasmid was linearized with the restriction enzyme, NotI and transformed with standard electroporation transformation and selected with EMM-leu media. The deletions of various genes were made with either PCR-based gene-targeting approaches or by a cross followed by random-spore analysis.

### Microscopy

Cells were grown to mid-log phase in YEA media at 25°C and imaged. Fluorescence images were acquired using a widefield epifluorescence microscope equipped with a 20×/0.8 NA oil immersion objective and a GFP filter set. Exposure times and acquisition settings were kept constant for all samples. Image processing was performed in Fiji (ImageJ). All image adjustments (brightness, contrast) were applied equally across the entire image and across samples for display purposes only.

### FACS analysis

Cells corresponding to different genotypes were harvested in mid-log phase. 3 x 10^7^ cells/ml were washed with water and then fixed with 70% ethanol/30% sorbitol. The cells were washed three times with 1X tris-buffered saline (TBS) and resuspended in 1mL 1xTBS. GFP fluorescence was measured using a Beckman Cytoflex 4 FACS analyzer with a wavelength of 488nm. A primary gate based on physical parameters (forward and side light scatter) was set to exclude dead cells or debris. 10,000 cells were analyzed for each sample.

### Silencing assays

Cells were grown in 3 mL of yeast extract containing adenine (YEA) at 32C overnight. Cells were washed twice in water and then resuspended to a concentration of 3 x 10^8^ cells/ml. 5uL of 10-fold serial dilutions was spotted on indicated plates (YE, YE + FOA, YE+tet, YE+FOA+tet). Plates were incubated for 2–5 days before photographing.

### Microfluidics

For single-cell analysis, cells were grown to mid-log phase (OD600 = 1.0) in YEA media at 32°C. Cells were then loaded into a custom-fabricated PDMS-based microfluidics channel. Fresh YEA media was perfused continuously at a flow rate of 25 µL/min, and the chip was maintained at 32°C throughout the experiment. Cells were imaged for about 2 days with snapshots of GFP emission captured every hour to reduce phototoxicity.

### RNA extraction

10mL of cells were grown to late log phase (OD600-1.8-2.2) in YEA medium. Cells were resuspended in 750µL TES buffer (0.01M Tris pH7.5, 0.01M EDTA, 0.5% SDS). Immediately 750µL of acidic phenol chloroform was added and vortexed for 2 min. Samples were incubated at 65°C for 40 min, vortexing for 20 sec every 10 min. The aqueous phase was separated by centrifuging in Phase Lock tubes for 5 min at 13000 rpm at 4°C. The aqueous phase was transferred to new tubes and ethanol precipitated. After extraction, RNA was treated with DNase. Then the RNA was cleaned up using RNeasy Mini kits (QIAGEN).

### RNA-seq analysis

polyA enriched libraries were prepared, and sequenced using services offered by Novogene. Raw FastQ files were aligned to the ASM294v2 reference genome using STAR and then indexed using samtools. Bam files were grouped by genotype replicate and differential expression analysis was performed through Defined Region Differential Seq in the USEQ program suite (Love et al., 2014).

### Chromatin immunoprecipitation (ChIP)

30 mL of cells were grown to late log phase (OD600-1.8-2.2) in yeast extract supplemented with adenine or yeast extract supplemented with adenine containing FOA (1g/L) and fixed with 1% formaldehyde for 15 min at room temperature (RT). 130mM glycine was then added to quench the reaction and incubated for 5 min at RT. The cells were harvested by centrifugation, and washed with TBS (50 mM Tris, pH 7.6, 500 mM NaCl). Cell pellets were resuspended in 300 µL lysis buffer (50 mM HEPES-KOH, pH 7.5, 100 mM NaCl, 1mM EDTA, 1% Triton X-100, 0.1% SDS, and protease inhibitors) to which 500µL 0.5 mm glass beads were added and cell lysis was carried out by bead beating using Omni Bead Ruptor at 3000 rpm for 30sec x 10 cycles. Tubes were punctured and the flowthrough was collected in a new tube by centrifugation which was subjected to sonication to obtain fragment sizes of roughly 100500 bp long. After sonication, the extract was centrifuged for 15 min at 13000 rpm at 4°C. The soluble chromatin was then transferred to a fresh tube and normalized for protein concentration by the Bradford assay. For each normalized sample, 25µL lysate was saved as input, to which 225µL of 1xTE/1% SDS were added (TE: 50 mM Tris pH 8.0, 1 mM EDTA). Protein A Dynabeads were preincubated with antibody. For each immunoprecipitation, 2 µg H3K9me2 antibody (ab1220, Abcam) or 2µg H3K9me3 antibody (39161, Active Motif) coupled to 30µL beads was added to 400µL soluble chromatin and the final volume of 500uL was achieved by adding lysis buffer.

Samples were incubated for 3h at 4°C. The beads were collected on magnetic stands and washed 3 times with 1 mL lysis buffer and once with 1 mL TE. For eluting bound chromatin, 100µL elution buffer I (50 mM Tris pH 8.0, 10mM EDTA, 1% SDS) was added, and the samples were incubated at 65°C for 5 min. The eluate was collected and incubated with 150 µL 1xTE/0.67% SDS in the same way. Input and immunoprecipitated samples were finally incubated overnight at 65°C to reverse crosslink. 60µg glycogen, 100 mg proteinase K (Roche), 44 ml of 5M LiCl, and 250 ml of 1xTE was added to each sample, and incubation was continued at 55C for 1h. Phenol/chloroform extraction was carried out for all the samples, followed by ethanol precipitation. Immuno-precipitated DNA was resuspended in 100µL of 10 mM Tris pH 7.5 and 50 mM NaCl. ChIP experiments were analyzed using quantitative PCR with Taq polymerase and SYBR Green using a CFX Opus 384 Real-Time PCR System.

### ChIP-seq library preparation and processing

Libraries were constructed using the manufacturer’s guidelines in the NEBNext Ultra II FS DNA Library Prep Kit for Illumina, using 1ng of starting material. Barcoded libraries were pooled and sequenced with next-generation sequencing. First, raw reads were demultiplexed by barcode.

Then the sequences were trimmed with Trimmomatic, aligned with BWA, and normalized by counts per million (Bolger et al., 2014; Li and Durbin, 2010). Then the reads were visualized with IGV. For further analysis peaks were called using MACS2 with -g 1.27e6 in broad mode with a cutoff of 0.05 (Y. Zhang et al., 2008). Heatmaps were generated using deepTools (v3.5.1) (Ramírez et al., 2016).

### Genome-wide screen

Genome-wide screen was performed as previously described with some modifications(Muhammad et al., 2024). A haploid deletion library (Bioneer, version 3.0) was crossed with strains containing *leu1+:TetR-Mit1* and *ade6+-10XtetO-ura4+* reporter genes at the *ura4+* locus by using RoToR HAD colony pinning robot (Singer Instruments). Following mating, cultures were incubated on SPAS plates at 42°C for 4 days. Spores were plated on plates lacking leucine (EMM-LEU), then dropout uracil plates (EMM-URA), before selection on YES containing G418. Cells were then transferred to YES (non-selective) and YES supplemented with 1 g/L 5’-fluororotic acid (EMM+FOA) for the readout. Colonies were photographed and sizes were measured using the Gitter R package (https://omarwagih.github.io/gitter/). To calculate relative growth (reporter activity) c colony sizes on YES+FOA were divided by growth on non-selective media (YES). All values were normalized to the median values of the individual 384-plates. Log2 values were used for clustering and data visualization. The deletion library was screened three times independently , each biological replicate containing two technical replicates from the same genetic cross generated by duplication during the germination step. Mating type locus data published previously was used as comparison for the newly generateddatasets (Muhammad et al., 2024).

### AlphaFold3 structure predictions

Protein complex structures were predicted using AlphaFold Server that uses AlphaFold3 for structure predictions(Abramson et al., 2024). Amino acid sequences of candidate interacting proteins were first analyzed using metapredict2 to identify intrinsically disordered regions (IDRs), which were removed to improve structural prediction (Lotthammer et al., 2024). Truncated sequences lacking predicted IDRs were submitted to AlphaFold Server using default settings, with the number of models set to five per run. Predicted complex structures were analyzed using AlphaBridge to examine interaction interfaces and residue–residue contact networks (Álvarez-Salmoral et al., 2025).

### Yeast Two Hybrid assays

PCR amplified sequences of SHREC and CLRC complex protein-pairs were cloned into the pGBDU-C1 (DNA-binding domain, BD) and pGAD-C1 (activation domain, AD) vectors, using Gibson cloning. Constructs were confirmed by whole plasmid sequencing. Plasmids were co-transformed into the Saccharomyces cerevisiae strain PJ69–4a (harboring selectable *GAL* UAS-dependent HIS3 and ADE2 reporter genes) using lithium acetate. Transformants were selected on synthetic dropout medium lacking leucine and tryptophan (SC-URA-LEU). Protein–protein interactions were assessed by spotting serial dilutions of co-transformants onto SC-LEU-URA-HIS plates. Growth and reporter activation were assessed after 3–5 days of incubation at 32°C.

---

**Figure S1.**
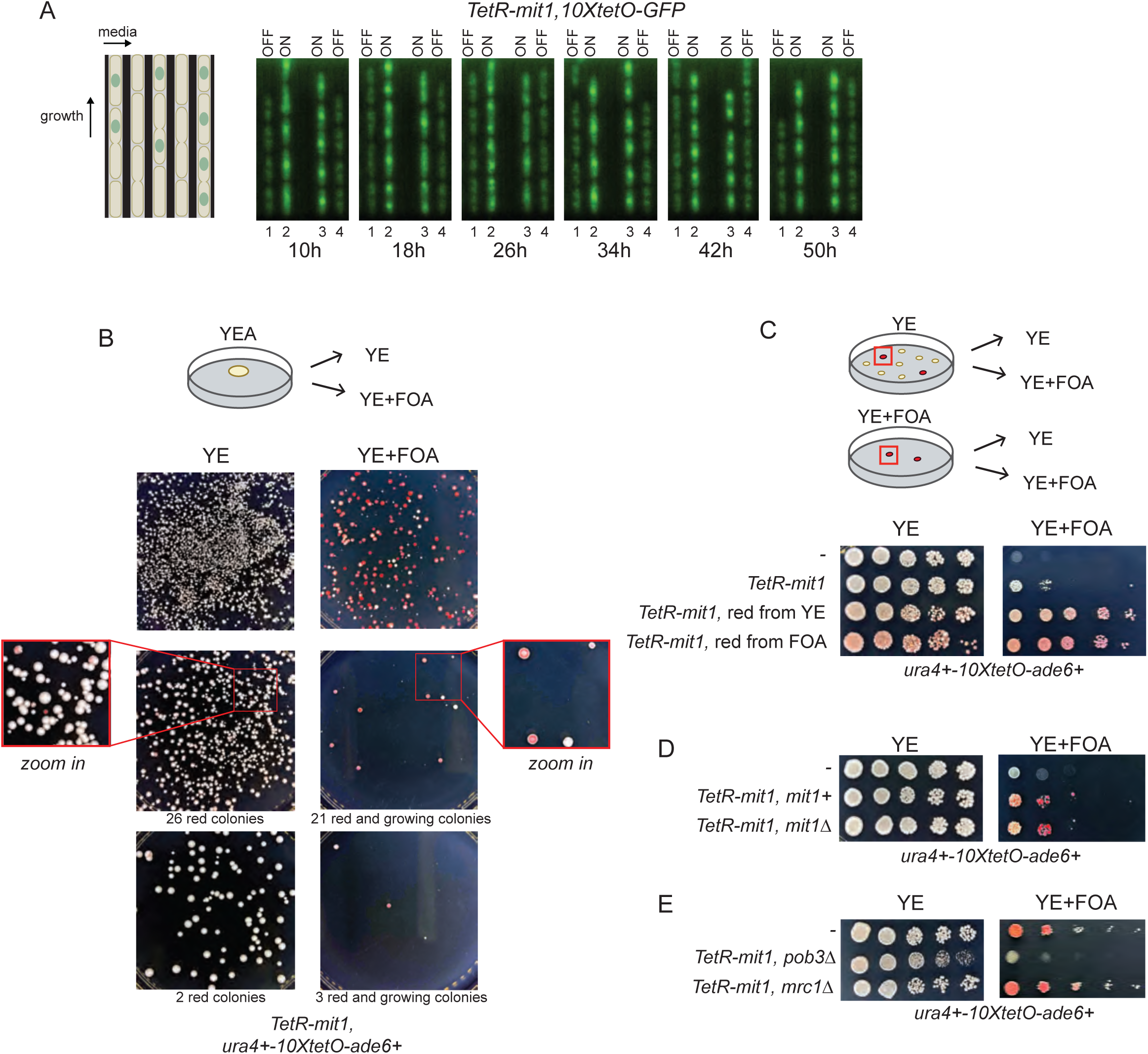
Lineage tracing measurements and plating assays measure establishment frequencies upon tethering Mit1. (A) A microfluidic device enables tracking of individual cell lineages for over 50 hours. Strains from **1C** were continuously imaged. Snapshots at 8-hour intervals are shown with each frame capturing four independent lineages. Cells are either GFP-ON or GFP-OFF. (B) Silencing assay of *TetR-mit1* on non-selective and FOA selective media. Cells are plated at ten-fold dilutions (top to bottom). (C) Silencing assay of *TetR-mit1* red colonies selected from either non-selective (YE) or selective (YE+FOA) plates in the *ura4+ – ade6+* reporter strain. (D) Silencing assay in strains expressing *TetR-mit1* with and without endogenous Mit1 (*mit1+* versus *mit1Δ)* in the *ura4+ – ade6+* reporter strain. (E) Silencing assay measuring silencing in the absence of histone chaperones, Pob3 and Mrc1. For all plate-based assays, cells are plated at 10-fold serial dilutions on non-selective low adenine media (YE) and on low adenine media containing FOA (YE+FOA).

**Figure S2.**
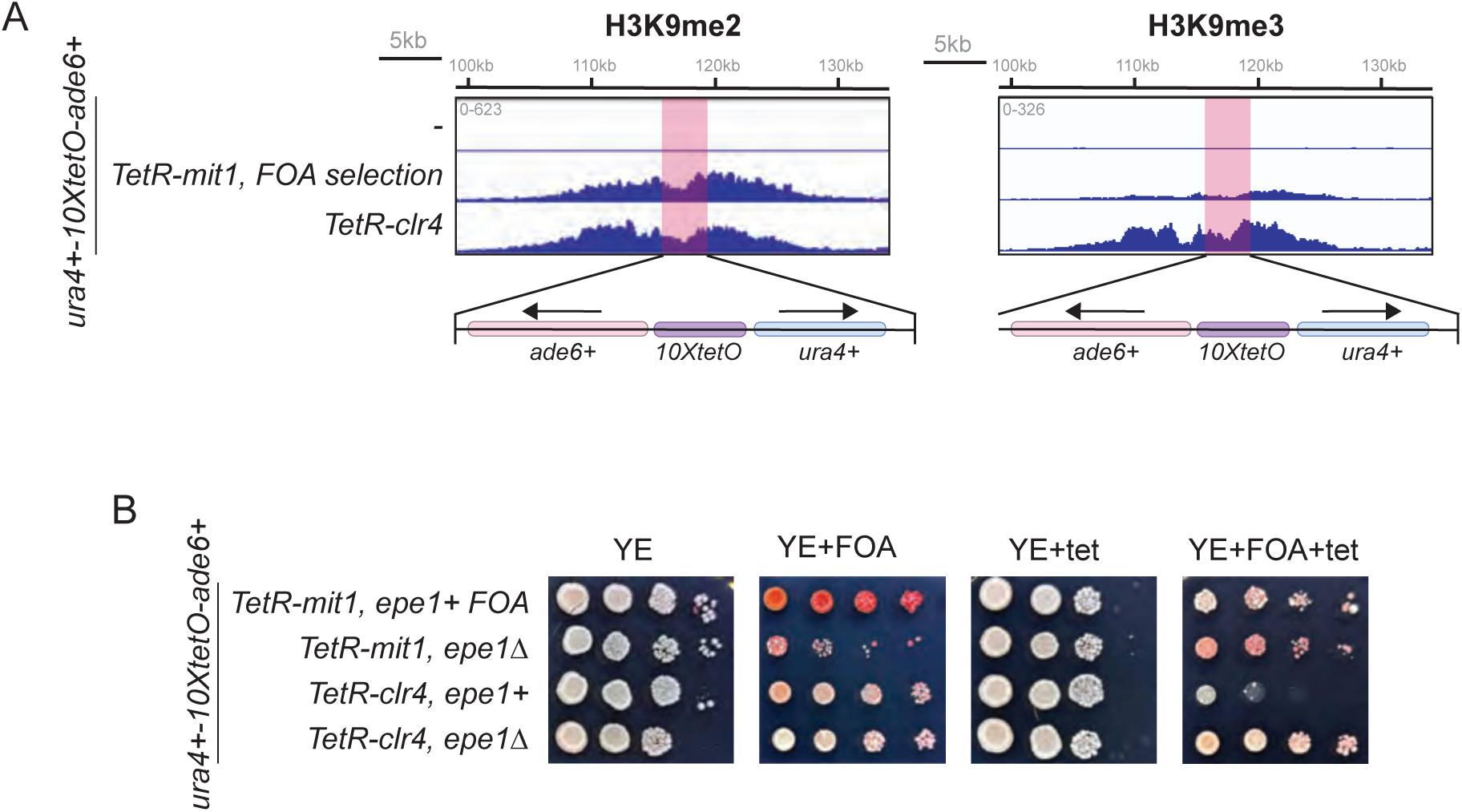
Comparison of Mit1-initiated versus Clr4-initiated silencing. (A) ChIP-seq of H3K9me2 *(same as 2*D) and H3K9me3 (*same as 2*E) flanking the *10xTetO* binding site in TetR-Mit1 versus TetR-Clr4 expressing strains. (B) Adding tetracycline leads to TetR dissociation enabling measurement of H3K9me maintenance independent of sequence-dependent establishment. Silencing assay comparing *epe1+* versus *epe1Δ* cells with TetR-Mit1 without or with tetracycline (+tet) in the *ura4+ – ade6+* reporter strain. Supplemental Figure 3: Systematic comparison of factors involved in mating type and Mit1-mediated heterochromatin

**Figure S3.**
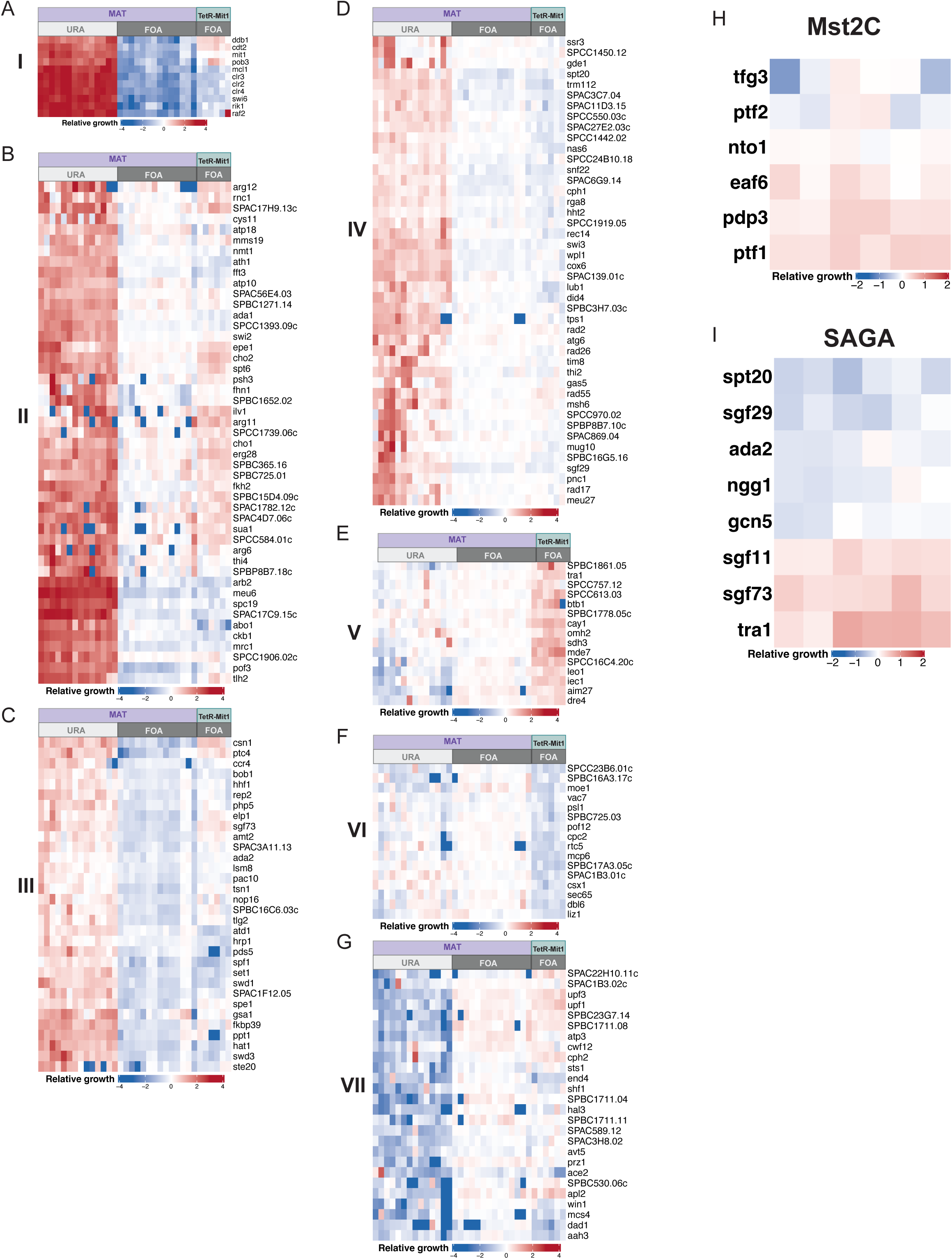
Systematic comparison of factors involved in mating type and Mit1-mediated heterochromatin. Heatmaps displaying clusters of deletions in both a *ura4+* reporter at the endogenous heterochromatin mat locus and at the *ade6+-ura4+* reporter locus in *TetR-mit1* cells. Relative growth values are log2-transformed. **(A)** Heatmap (Cluster-I) displaying comparisons of reporter activity between core-heterochromatin protein deletions in a *ura4+* reporter at the endogenous heterochromatin *mat* locus and in the *ade6+-ura4+* reporter in *TetR-mit1* cells. (**B, C**) Heatmap (Cluster-II and III) shows mutants with moderate defects at the mating type locus but mixed silencing and anti-silencing effects in the Mit1 system. (**D**) Heatmap (Cluster-IV) shows mutants with uniformly weak silencing defects in both mating type locus and Mit1 systems. (**E** Heatmap (Cluster-V) shows an enrichment of mutants that display improved silencing with a heightened sensitivity in Mit1-mediated repression. (F) Heatmap (Cluster-VI) shows mutants that uniquely affect Mit1-dependent silencing. (G) Heatmap (Cluster-VII) shows mutants that improve both mating type silencing and Mit1-mediated repression. (H) Heatmap displaying relative growth scores for Mst2C deletions in *TetR-mit1* strains. (I) Heatmap displaying relative growth scores for SAGA complexes deletions in *TetR-mit1* strains.

**Figure S4.**
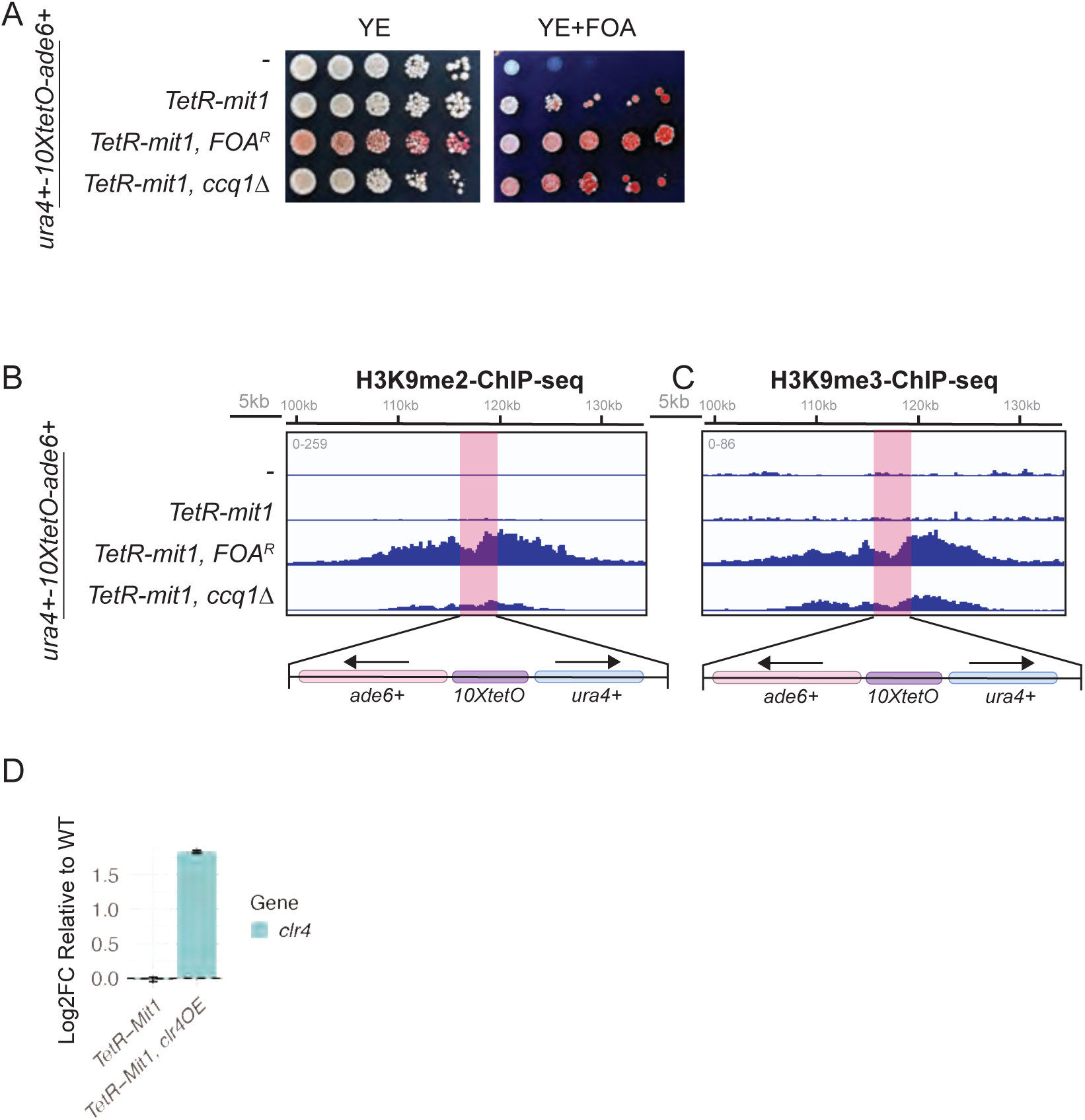
Increasing the dosage of heterochromatin factors by disrupting endogenous heterochromatin domains enhances Mit1-initiated silencing. (A) Silencing assay for *TetR-mit1* expressing cells in a *ccq1Δ* background. (**B, C**) ChIP-seq of H3K9me2 (**B**) and H3K9me3 (**C**) flanking the TetR-Mit1 binding site in the indicated genotypes. The *ura4+ – ade6* reporter which replaces the endogenous *ura4* locus is highlighted in red. A 50kb region spanning the tethering site is shown. (**D**) RNA-seq data examining expression levels of *clr4* upon overexpression. Changes in expression are shown as log2 fold change normalized to wild-type cells (N=3). All silencing assays were performed using the ura4+ – ade6+ reporter strain.

**Figure S5.**
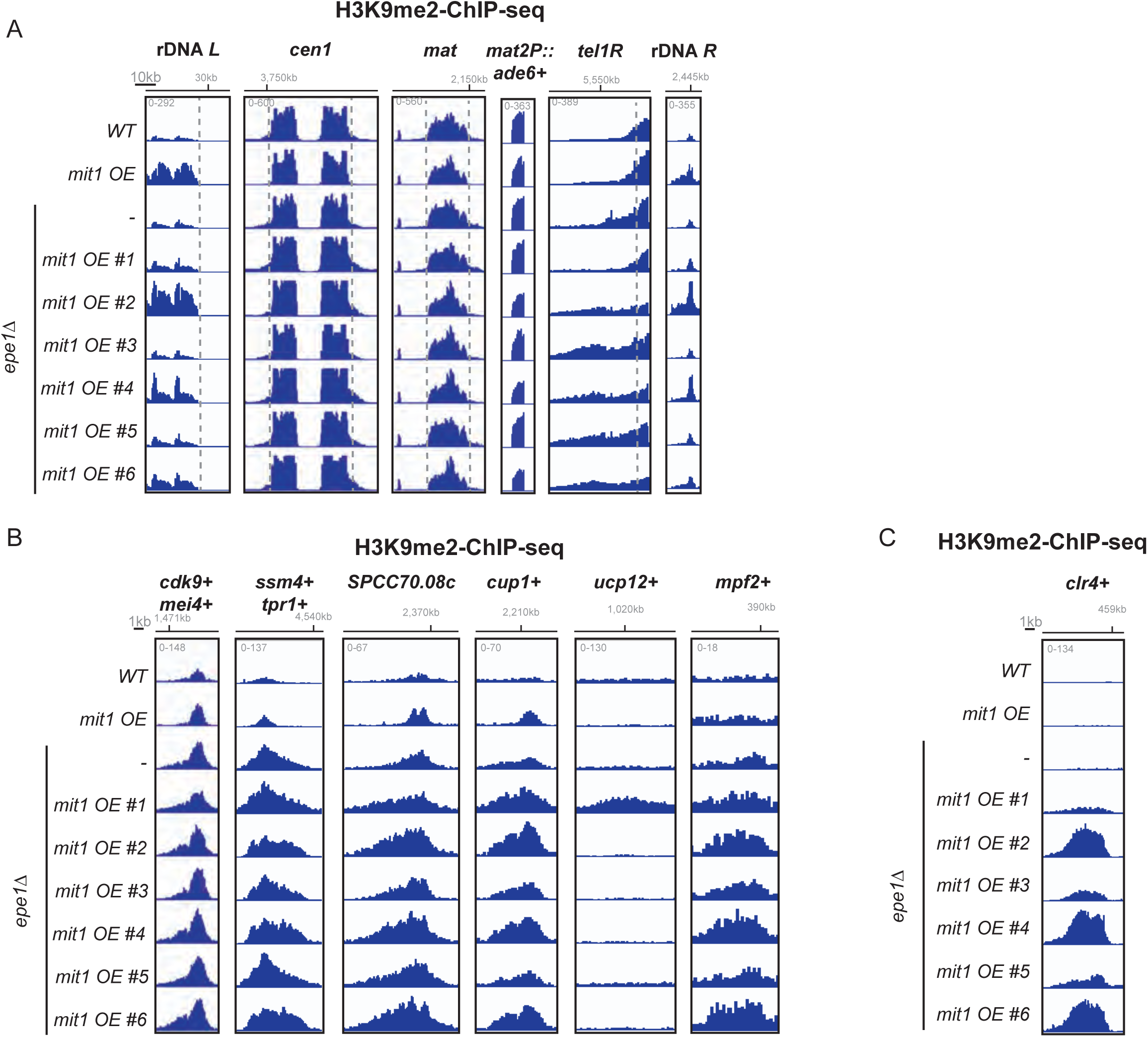
Mit1 overexpression leads to H3K9me2 spreading at constitutive heterochromatin and increased H3K9me2 deposition at heterochromatin islands. (A) ChIP-seq of H3K9me2 at endogenous heterochromatin regions in the indicated genotypes. (B) ChIP-seq of H3K9me2 at regions with *de novo* peaks in *Mit1 OE*. (C) ChIP-seq of H3K9me2 at the *clr4* locus.

